# Genetic context modulates aging and degeneration in the murine retina

**DOI:** 10.1101/2024.04.16.589625

**Authors:** Olivia J. Marola, Michael MacLean, Travis L. Cossette, Cory A. Diemler, Amanda A. Hewes, Alaina M. Reagan, Daniel A. Skelly, Gareth R. Howell

## Abstract

**Background:** Age is the principal risk factor for neurodegeneration in both the retina and brain. The retina and brain share many biological properties; thus, insights into retinal aging and degeneration may shed light onto similar processes in the brain. Genetic makeup strongly influences susceptibility to age-related retinal disease. However, studies investigating retinal aging have not sufficiently accounted for genetic diversity. Therefore, examining molecular aging in the retina across different genetic backgrounds will enhance our understanding of human-relevant aging and degeneration in both the retina and brain—potentially improving therapeutic approaches to these debilitating conditions.

**Methods:** Transcriptomics and proteomics were employed to elucidate retinal aging signatures in nine genetically diverse mouse strains (C57BL/6J, 129S1/SvlmJ, NZO/HlLtJ, WSB/EiJ, CAST/EiJ, PWK/PhK, NOD/ShiLtJ, A/J, and BALB/cJ) across lifespan. These data predicted human disease-relevant changes in WSB and NZO strains. Accordingly, B6, WSB and NZO mice were subjected to human-relevant *in vivo* examinations at 4, 8, 12, and/or 18M, including: slit lamp, fundus imaging, optical coherence tomography, fluorescein angiography, and pattern/full-field electroretinography. Retinal morphology, vascular structure, and cell counts were assessed *ex vivo*.

**Results:** We identified common molecular aging signatures across the nine mouse strains, which included genes associated with photoreceptor function and immune activation. Genetic background strongly modulated these aging signatures. Analysis of cell type-specific marker genes predicted age-related loss of photoreceptors and retinal ganglion cells (RGCs) in WSB and NZO, respectively. Fundus exams revealed retinitis pigmentosa-relevant pigmentary abnormalities in WSB retinas and diabetic retinopathy (DR)-relevant cotton wool spots and exudates in NZO retinas. Profound photoreceptor dysfunction and loss were confirmed in WSB. Molecular analyses indicated changes in photoreceptor-specific proteins prior to loss, suggesting photoreceptor-intrinsic dysfunction in WSB. In addition, age-associated RGC dysfunction, loss, and concomitant microvascular dysfunction was observed in NZO mice. Proteomic analyses revealed an early reduction in protective antioxidant processes, which may underlie increased susceptibility to DR-relevant pathology in NZO.

**Conclusions:** Genetic context is a strong determinant of retinal aging, and our multi-omics resource can aid in understanding age-related diseases of the eye and brain. Our investigations identified and validated WSB and NZO mice as improved preclinical models relevant to common retinal neurodegenerative diseases.

## Background

Retinal degeneration is a leading cause of vision loss and is among the most common forms of neurodegenerative disease. These conditions, including age-related macular degeneration (AMD), diabetic retinopathy (DR), glaucoma, and retinitis pigmentosa (RP), pose a major economic burden (1), and loss of vision due to these conditions severely reduces quality of life (2-4). Aging is one of the most important risk factors for the development of retinal degenerative diseases (5, 6). Given the shared biological features of the retina and brain, mechanisms driving retinal degenerative diseases are likely relevant to neurodegenerative diseases of the brain, such as Alzheimer’s disease and related dementias. Furthermore, there is growing interest in leveraging the accessibility of the retina to track the progression of neurodegeneration within the brain (7-9). Yet, age-related and pathological changes in retinal function may obscure this insight into brain pathology. Thus, it is vital to understand the mechanisms of healthy and pathological retinal aging.

Only a few studies have investigated aging-associated transcriptomic changes of the human retina. One such study showed evidence of changes in cellular metabolism, cell cycle regulation, and cell adhesion in aged retinas compared to young controls (10). Mouse models have also aided in understanding retinal aging and degeneration (11, 12). Previous retinal microarray analyses revealed aging-associated complement activation concomitant with changes in stress response pathways in humans and C57BL/6J (B6) mice (13, 14). B6 rod photoreceptors exhibit age-associated changes to metabolic pathways—including lipid metabolism (15, 16). These initial studies suggest human and mouse retinas exhibit similarities in aging responses. The development of higher throughput transcriptomics and proteomics has made it possible to profile many more transcripts and proteins in a single sample (17). This is critically important for retinal tissue, in which rods comprise ∼70% of cells (18, 19). In-depth profiling of the retina transcriptome and proteome may reveal changes that are smaller in magnitude or are driven by changes in less abundant cell types—identifying novel therapeutic targets to explore. There is a substantial genetic component influencing susceptibility to retinal degeneration. For instance, there are growing numbers of single nucleotide polymorphisms linked to retinal neurodegenerative diseases (20-23). In addition, genetic contributions to retinal degeneration are likely to be underestimated as, historically, the inclusion of genetically diverse individuals is lacking (24). One of the only studies investigating aging in the human retina was limited to Caucasians, and another study did not report ethnicities (10, 14). While there is a similar gap in preclinical studies of retinal neurodegenerative diseases (the majority incorporate only one strain– commonly B6), a few studies show genetic background strongly influences retinal disease phenotypes. For example, in retinal degeneration, the *rd7* phenotypes on a B6 background were suppressed by protective alleles present in CAST and AKR/J genetic backgrounds (25). Furthermore, a quantitative trait locus mapping study identified multiple loci that modified *rd3*-associated retinal phenotypes (26). For glaucoma, mutations known to drive ocular hypertension and consequential retinal ganglion cell loss (RGCs) in DBA/2J mice, failed to cause glaucoma-relevant phenotypes in B6 mice (11). Thus, the inclusion of genetically diverse mouse strains will be critical in the study of retinal aging and degeneration.

Mouse resource populations consisting of genetically diverse strains or individuals (e.g. Collaborative Cross (CC) and Diversity Outbred (DO) populations) have been leveraged to link genetic context to specific phenotypes and gene expression (27-30). CC and DO mice originate from crosses of eight founder strains, which encapsulate the majority of known genetic differences within laboratory mice (27, 28). Aging has been linked to numerous cellular perturbations across many tissues (31, 32). We have documented the importance of incorporating diverse mouse strains into studies of aging and neurodegenerative diseases of the brain (33-37), yet their use in understanding retinal aging has been limited. Therefore, we profiled molecular signatures of retinal aging across nine genetically diverse mouse strains: standard pigmented strains (B6, 129S1, NZO), standard albino strains (BALBc, NOD, AJ), and wild-derived strains (WSB, CAST, PWK). These strains include all founder strains of the CC and DO mouse panels with the addition of commonly utilized BALBc mice. Our initial analyses of these data unveiled a common aging signature among mice that was strongly modulated by genetic background. Analyses of cell type-specific marker genes predicted NZO and WSB strains may exhibit human-relevant retinal neurodegeneration with age. *In vivo* and *ex vivo* ophthalmologic assays indicated hallmarks of RP- and AMD-relevant photoreceptor degeneration in WSB retinas, while NZO mice displayed many features relevant to DR. Collectively, our work elucidates the mechanisms underlying retinal aging and degeneration and offers a publicly available database to accelerate research into mechanisms relevant to aging and degeneration of both the retina and brain.

## Methods

### Mouse strains and cohort generation

All research was approved by the Institutional Animal Care and Use Committee (IACUC) at The Jackson Laboratory. All mice were bred and housed in a 12-hour light/dark cycle and received a standard 6% LabDiet Chow (Cat# 5K52) and water *ad libitum*. Cohorts of the following mouse strains were established and aged together for multi-omics analyses: C57BL/6J (B6; JAX Stock# 000664), 129S1/SvlmJ (129S1; JAX Stock# 002488), NZO/HlLtJ (NZO; JAX Stock# 002105), WSB/EiJ (WSB; JAX Stock# 001145), CAST/EiJ (CAST; JAX Stock# 000928), PWK/PhJ (PWK; JAX Stock# 003715), A/J (AJ; JAX Stock# 000646), NOD/ShiLtj (NOD; JAX Stock# 001976), BALB/cJ (BALBc; JAX Stock# 000651). Additional cohorts of B6, NZO, and WSB mice were generated for all other experiments.

### Molecular Investigations

### Tissue collection for RNA-sequencing and Proteomics

Mice were anesthetized with ketamine/xylazine and then underwent cardiac perfusion with PBS. Eye globes were gently excised, and retinas dissected. Retinas were quickly minced with a fresh razor blade on ice and tissue dispersed between two DNAse/RNAse free microcentrifuge tubes. Samples were flash frozen in liquid nitrogen and stored at -80°C until further processing.

### RNA Isolation and RNA-sequencing

Total RNA was isolated from retinal tissue aliquots using RNeasy Micro kits (Qiagen), according to the manufacturers’ protocols, including the optional DNase digest step. Tissues were homogenized in RLT buffer (Qiagen) using a Pellet Pestle Motor (Kimbal). RNA concentration and quality were assessed using the Nanodrop 2000 spectrophotometer (Thermo Scientific) and the RNA 6000 Nano and Pico Bioanalyzer Assay (Agilent Technologies). Libraries were constructed using the KAPA mRNA HyperPrep Kit (Roche Sequencing and Life Science), according to the manufacturer’s protocol. Libraries were assessed using D5000 ScreenTape (Agilent Technologies) and the Qubit dsDNA HS Assay (ThermoFisher) according to the manufacturers’ instructions. Approximately, 44M 150bp paired-end reads were sequenced per sample on an Illumina NovaSeq 6000 using the S4 Reagent Kit v1.5 (Illumina, Cat#20028313) by the Genome Technologies Core at The Jackson Laboratory.

Raw FASTQ files were processed using standard quality control practices. High quality read pairs were aligned to a custom pseudogenome produced by incorporating strain-specific variants (single nucleotide variants and short insertions/deletions derived from Mouse Genomes Project REL-1505)(38) into the mouse reference genome (GRCm38/mm10) using g2gtools version 0.2.7 (https://github.com/churchill-lab/g2gtools). Reads were aligned using STAR version 2.7.10a. Transcript counts were generated using RSEM version 1.3.3 (39).

Differential expression analyses were completed using edgeR v3.40.2 within the R environment v4.2.3 using the glmQLFtest function (40). filterByExpr within edgeR was used to filter our very lowly expressed genes prior to TMM normalization (40). Differential expression was analyzed using generalized linear models via edgeR, with either 1) groups defined as a combination of strain, sex, age and a library batch term (∼0+Group+Batch) and contrasts designed to test for average strain effect, average age effect within each strain; or 2) age as a continuous variable to control and test for differential expression related to strain differences or determining the impact of aging while controlling for strain and sex differences. The clusterProfiler v4.6.2 R package was used to test for overrepresentation of gene ontology (GO) Biological Process gene sets within the list of differentially expressed genes (DEGs) with a false-discovery rate (FDR) of less than 0.05 and enrichPlot v1.18.4 and EnhancedVolcano v1.16.0 were used to visualize the enrichment results (41-47).

### Proteomics

#### Sample Preparation

Each half retina sample had ice-cold 600 µL of extraction buffer (2:2:1 methanol:acetonitrile:water) added, along with a pre-chilled 5 mm stainless steel bead (QIAGEN). Tubes were then added to the pre-chilled (at -20°C) cassettes for the Tissue Lyser II in batches of 48 samples, then were lysed in the Tissue Lyser II for 2 minutes at 30 1/s. The stainless steel beads were removed with a magnet and the samples were placed at -20°C for 16h overnight. Sample extracts were then centrifuged at 21,000 x g at 4°C for 15 minutes. The supernatant was removed, and the protein pellet was reconstituted in 100 µL of 50 mM HEPES buffer, pH 8.0. Reconstituted samples were vortexed at max speed for 30 seconds, a chilled steel bead was added, and they were reconstituted using the Tissue Lyser II as described for the extraction. Samples were then waterbath sonicated (sweep at 37 Hz at 100% power) for 5 minutes (30 seconds on, 30 seconds off for five cycles) with ice added to keep the temperature from rising. Samples were assessed for clarity and if necessary, the sonication was repeated. Samples were then centrifuged at 21,000 x g for 15 minutes at 4°C and protein supernatant was transferred to a clean microcentrifuge tube. Protein quantification was then performed using a microBCA assay (Thermo, Cat.# 23235) on a single well for each sample with a 1:25 dilution. The remaining sample was snap-frozen and stored at -80°C until further use.

#### Protein Digestion and Peptide Purification

After protein quantification, 20 µg of each sample were aliquoted to a new tube and brought to an equal volume in 50 mM HEPES, pH 8.0. Samples with less than 20 ug total required use of the full sample for digest and the trypsin ratio was adjusted at the digestion stage. Samples were reduced with 10 mM dithiothreitol at 42°C for 30 minutes on a ThermoMixer with 500 rpm agitation, alkylated with 15 mM iodoacetamide for 20 minutes at room temperature on a ThermoMixer with 500 rpm agitation, and trypsin-digested (Sequence Grade Modified; Promega) with a 1:50 trypsin:protein ratio at 37°C for 20h. Following the digest, all peptide samples were C18-purified using Millipore C18 zip-tips (Millipore, Cat#: ZTC18S096) according to the manufacturer protocol’s and as previously reported (48). Eluted purified peptides were dried with a vacuum centrifuge and stored at -20°C until tandem mass tag (TMT) labeling was performed.

#### TMT Pro Peptide Labeling

The dried peptide samples were reconstituted in 20 uL of 50 mM HEPES, pH 8.5) and quantified using the Quantitative Colorimetric Peptide Assay (Thermo, Cat# 23275) according to the manufacturer’s protocol. Samples were diluted to a concentration of ∼0.5 µg/µL in 10 µL of 50 mM HEPES, pH 8.5, and mixed on a ThermoMixer for 10 minutes at 25°C (500rpm). To create a quality control pool for a carrier channel, equal amounts of all samples were used to create enough pool that could be used in each multiplex for normalization. While the peptide solutions mixed, TMTpro reagents (Thermo, Cat# A44520) were mixed according to manufacturer protocol. TMTpro labels were added to a 19.37 mM final concentration to the appropriate peptide sample, followed by a 1hr incubation at 25°C with 500 rpm agitation on a ThermoMixer. As the study required many multiplexes the samples were first randomized for the multiplex assignment and then were randomized for the TMT tag assignment within a multiplex through a list randomization in Random.org. Reactions were then quenched by adding a final concentration of 0.5% hydroxylamine and incubating for 15 minutes at 25°C with 400 rpm agitation on a ThermoMixer. All samples were added to the appropriate TMT multiplex group, acidified by adding 20 uL of 10% formic acid, snap-frozen, and stored at -80°C overnight. The next day samples were dried in a vacuum centrifuge.

#### Liquid Chromatography Tandem Mass Spectrometry Analysis (LC-MS/MS)

Each of the dried multiplexed peptide samples were reconstituted in 25 uL of LC-MS grade water with 0.1% TFA and zip-tipped using Millipore C18 zip-tips according to the manufacturer protocol’s and as previously reported (48). Purified multiplexes were then dried in a vacuum centrifuge and reconstituted in 20 µL of 98% H2O/2% ACN with 0.1% formic acid via pipetting and vortexing. Reconstituted samples were then transferred to a mass spec vial and placed in the autosampler at 4°C. LC-MS/MS was then performed on a Thermo Eclipse Tribrid Orbitrap with a FAIMS coupled to an UltiMate 3000 nano-flow LC system in The Jackson Laboratory Mass Spectrometry and Protein Chemistry Service. The method duration was 180 minutes at a flow rate of 300 nL/min. Buffer A (100% H2O with 0.1% formic acid) and Buffer B (100% acetonitrile with 0.1% formic acid) were utilized for the gradient. The full gradient consisted of 98% A/2% B from 0-2 minutes, 98% A/2% B at 2 minutes to 92.5% A/7.5% B at 15 minutes, 92.5% A/7.5% B at 15 minutes to 70% A/30% B at 145 minutes, and 70% A/30% B at 145 minutes to 10% A/90% B at 155 minutes. The gradient was held at 10% A/90% B until 160 minutes and brought to 98% A/2% B by 162 minutes, where it was then held for 15 minutes to equilibrate the column. The TMT SP3 MS3 real-time search (RTS) method was used on the instrument. Global parameters included a default charge = 2, expected peak width = 30 seconds, advanced peak determination, spray voltage = 2000V, mode = positive, FAIMS carrier gas =4.6 L/min, and an ion transfer tube temperature = 325°C. The instrument method utilized the FAIMS voltages of -40V, -55V, and -65V. Settings for precursor spectra detection (MS1) in each node included: cycle time = 1 second (each node), detector = Orbitrap, Orbitrap resolution = 120,000, scan range = 400-1600 m/z, RF lens % = 30, normalize AGC target (%) = 250, maximum inject time (ms) = auto, microscans = 1, data type = profile, polarity = positive, monoisotopic precursor selection = peptide, minimum intensity threshold = 5.0e3 (lower because of the FAIMS), charge states = 2-7, and a dynamic exclusion of a n = 1 for 60 seconds. Peptide fragment analysis (MS2) was performed in the ion trap and settings included: isolation window (m/z) = 0.7, collision energy (%) = 35 (fixed), activation type = CID, CID activation time = 10 ms, quadrupole isolation mode, ion trap scan rate = turbo, maximum inject time = 35 ms, and data type = centroid. Prior to data-dependent MS3 (ddMS3) analysis, RTS was utilized with the UniProtKB Mus musculus (sp_tr_incl_isoforms TaxID=10090) protein database including cysteine carbamidomethylation (+57.0215 Da), TMTpro16plex (+304.20171 on Kn), and methionine oxidation (+15.9949). Additional parameters in the RTS included maximum missed cleavages = 2, Xcorr threshold of 1, dCn threshold of 0.1, and a precursor ppm of 10. SP3 MS3 was performed in the Orbitrap and settings included SPS precursors = 20, isolation window = 0.7 m/z, activation type = HCD, HCD energy normalized at 45%, resolution = 60,000, scan range = 100-500 m/z, normalized AGC target = 500%, maximum injection time = 118 ms, and centroid data collection.

#### LC-MS/MS Data Analysis

All of the Thermo Eclipse RAW data files from the Eclipse Tribrid Orbitrap mass spectrometer were searched against the UniProtKB Mus musculus (sp_tr_incl_isoforms TaxID=10090) protein database in Proteome Discoverer (Thermo Scientific, version 2.5.0.400) using Sequest HT according to standard manufacturer recommended workflows. The Sequest protein database search parameters included trypsin cleavage, precursor mass tolerance = 20 ppm, fragment mass tolerance = 0.5 Da, static cysteine carbamidomethylation (+57.021 Da), static TMTpro modification on any N-terminus (+304.207 Da), and a variable methionine oxidation (+15.995 Da). Other setting included a maximum number of missed cleavages = 2, minimum peptide length = 6, a maximum of 144 amino acids, and a fragment mass tolerance = 0.6 Da. Percolator was used and in this module in the software, the target/decoy selection was concatenated, q-value validation was utilized, a maximum delta Cn = 0.05 was set, and FDR < 0.05 for all matches was used as the threshold. Default Minora node parameters were used. Abundance values were then normalized to the total ion signal in the samples using the Reporter Ions Quantifier Node for TMT-based MS3 events.

#### Differential Proteomics Analyses

Identified peptides with FDR<0.05 confidence and detected in at least three samples were used for further investigation. log_2_(scaled abundance+1) values were used for differential expression analyses. Differentially expressed proteins (DEPs) were determined using generalized linear models and a repeated measures model via limma and DEqMS (49, 50), which takes into account the number of unique peptide spectrum matches (PSMs) used for each peptide to enhance the statistical power of differential expression, with either 1) groups defined as a combination of strain, sex, age and a TMT multiplex batch term (∼0+Group+Multiplex) and contrasts designed to test for strain effects, age effects within each strain; or 2) age as a continuous variable to control and test for differential expression related to strain differences or determining the impact of aging while controlling for strain and sex differences. As samples were run in triplicate, we utilized the duplicateCorrelation function within limma to block on “mouseID”(50). As for the transcriptomic analyses, clusterProfiler v4.6.2 R package was used to test for overrepresentation of Gene Ontology (GO) Biological Process gene sets within the list of differentially expressed proteins with a FDR of less than 0.05 and enrichPlot v1.18.4 and EnhancedVolcano v1.16.0 were used to visualize the enrichment results (41-47). STRING-dB v 2.14.3 R package was used to visualize protein-protein interaction networks with the settings “score_threshold=200” and “network_type=physical” and to calculate enrichment of GO gene sets within the list of DEPs identified between 4M NZO or WSB with the other 5 pigmented strains (42, 51).

### In vivo ophthalmological investigations

All *in vivo* investigations were performed in B6, NZO, and WSB mice at 4, 8, 12, and/or 18M. *In vivo* techniques included: slit lamp imaging, fundus exams, fluorescein angiography, and electroretinography (full-field and pattern).

### Slit lamp imaging

Mice were anesthetized with ketamine/xylazine. Animals were placed on the apparatus with eyes aligned to the camera. Bright light images were obtained using Topcon DC-4 digital slit lamp and EZ Capture image software.

### Electroretinography (ERG)

Mice were placed in an anesthesia induction chamber infused with 3-4% isoflurane. Once fully anesthetized, mice were transferred to the heated platform of the Celeris-Diagnosys ERG system (Diagnosys LLC, MA, USA) to maintain body temperature at approximately 37°C, where anesthesia was sustained using 1-2% isoflurane. 1% Tropicamide and 2.5% phenylephrine were topically applied to dilate the pupils of both eyes, while a thin layer of 2.5% hypromellose was used to keep the eyes moist. Two corneal electrodes with integrated stimulators were placed on the surfaces of lubricated corneas to facilitate full-field ERG recordings. The Scotopic ERG was conducted initially, with nine steps of stimulus intensity (0.001, 0.002, 0.01, 0.0316, 0.316, 1,3, 10, and 31.6cd.s/m²). The final ERG waveform was the average of 10 individual waveforms. Following a 10-minute light adaptation interval, the photopic ERG was evoked by five steps of stimulus intensity (0.316, 1, 3, 10, 31.6cd.s/m²). The final ERG waveform was an average of 20 individual waveforms. The amplitude of the a-wave was measured as the difference between the pre-stimulus baseline and the trough of the a-wave, and the b-wave amplitude was measured from the trough of the a-wave to the peak of the b-wave. OP amplitudes were measured from the pre-stimulus baseline to the highest OP peak. The highest amplitude values across strains were elicited by 10cd.s/m^2^ luminance, thus, amplitudes generated at 10cd.s/m^2^ were compared.

### Pattern electroretinography (PERG)

All PERGs were performed using a JORVEC PERG system (Intelligent Hearing Systems, Miami, Florida) as described by the manufacturer. Briefly, mice were anesthetized with ketamine/xylazine and slit lamp was used to inspect each mouse eye prior to starting experiment. Mice were kept on a warming stage throughout the experiment. Electrodes were placed subcutaneously such that the active electrode is placed between the eyes with the tip just before the snout and the reference electrode is placed in line with the active electrode just between the ears. The ground electrode was placed approximately 2cm in front of the base of the tail. Scans were collected and waveforms were averaged using default settings. PERG amplitude was calculated as P1-N2 for each eye per mouse.

### Optical Coherence Tomography (OCT), fundus, and fluorescein angiography

OCT, fundus exams, and fluorescein angiographies were performed using Micron IV and Reveal OCT and Discover 2.4 software (Phoenix-Micron, Bend, OR). Mice were given 1 drop (20-30ul) of 0.5% Tropicamide (Somerset Therapeutics, NDC# 70069-121-01) in both eyes. After 10min, one drop (20-30ul) of 2.5% phenylephrine (Bausch & Lomb, NDC# 42702-0103-05) was applied to both eyes. Mice were then anesthetized using 4% isoflurane until a proper plane of anesthesia is achieved. Mice were then transferred to a nose cone on a heated holding cradle. Anesthesia was then reduced to 2% isoflurane. Eyes were kept moist using GenTeal Tears Lubricant Eye Gel Drops (Alcon). Respiration rate was monitored, and the camera lens was adjusted to be perpendicular to the cornea.

For OCT imaging, focus and brightness of the image guided OCT was adjusted until optimal image is previewed. OCT images were captured from the temporal to the nasal retina in a plane that included the optic nerve. To improve resolution, 10-40 images were taken and averaged. For fundus exams, a white light fundus image was taken of each eye. Brightness adjustments were made, and the focal plane was optimally adjusted. Immediately following the image-guided OCT and fundus imaging, mice were injected intraperitoneally with 1% Fluorescite® Fluorescein Sodium at a dose of 10mg/kg (10mg/mL solution, Akorn). Using a GFP filter, images of the right eye were taken every 30 seconds for 6 minutes. Images were taken of the left eye 30 seconds after the right eye imaging period.

Retinal layers from OCT images were measured using the FIJI distribution of ImageJ (52, 53). Each layer was measured 200, 400, and 600µm from the edge of the optic nerve using the line measurement tool, and the three respective measurements were averaged and used for statistical analyses.

### Fluorescein isothiocyanate–dextran (FITC-dextran) Leakage Assay

Mice were anesthetized with tribromoethanol and then transcardially perfused with 4% paraformaldehyde (PFA) (Electron Microscopy Services, Cat#15714) and 3mg/mL 70kDa fluorescein isothiocyanate–dextran (Millipore-Sigma, Cat#FD70S) in PBS. Eye globes were excised, and retinas were immediately dissected and flat mounted. Flat mounts were imaged with a Leica DMi8 inverted microscope within 30 minutes of harvest using a GFP filter by taking 10µm step z-stacked imaged to encompass the entire tissue at 10x magnification. The presence of leaks and avascular areas were recorded. Tissue integrity was assessed by bright field microscopy.

### Histological Analyses

At the time of sacrifice, animals were anesthetized with tribromoethanol and were transcardially perfused with PBS. Eyes were enucleated and fixed in 37.5% methanol and 12.5% acetic acid in PBS for 18h at 4°C. The Histology Core at The Jackson Laboratory performed paraffin embedding, sectioning, and hematoxylin and eosin and Prussian blue staining of optic nerve head-containing sections. Slides containing 2-3 sections per eye were imaged using a Hamamatsu S210 NanoZoomer Digital Slide Scanner at 40X magnification by the Microscopy Core at The Jackson Laboratory. Images were processed using custom FIJI scripts. Briefly, 400µm x 400µm regions of interest (ROIs) were selected on both sides of the optic nerve head at approximate distances of 500, 1000 and 1500µm representing the central, middle, and peripheral retina. After which, additional ROIs of the ONL per image were generated for automated thresholding, watershed segmentation, and quantification. The default thresholding was utilized with a size threshold of 30 for automated counting for all ONL cells.

### Vascular network isolation, staining, and analysis

At the time of sacrifice, animals were anesthetized with tribromoethanol and were transcardially perfused with PBS. Eyes were enucleated and fixed in 4% PFA in PBS for 18h at 4°C. The retinal vascular network was isolated as previously described (54) with a few modifications. Briefly, retinas were gently dissected and washed 3 times with ddH20 for 5minutes and then left overnight in ddH20 at RT. The following day, retinas were incubated in 2.5% Trypsin (ThermoFisher Scientific, Cat#15090046) for 2 hours at 37°C. After this, the retinas were washed 5 times with ddH20. Next, the vessels were gently washed from the neural retina with additional ddH20. Retinal vascular networks were then mounted onto glass slides and allowed to dry before staining. The vascular networks were stained with hematoxylin and eosin according to the manufacturer’s instructions (Abcam, Cat#ab245880). Stained networks were dehydrated with 3 washes with reagent alcohol and then mounted using organic mounting medium (Organo/Limonene Mount, Sigma-Aldrich, Cat#O8015). Dried slides were imaged using a Hamamatsu S210 NanoZoomer Digital Slide Scanner at 40X magnification by the Microscopy Core at The Jackson Laboratory. Acellular capillaries were manually counted across 4-7 400-500µm x 400-500µm ROIs of each retina, and the average count was used for statistical analyses.

### Immunohistochemistry (IHC) Analyses

At the time of sacrifice, animals were anesthetized with tribromoethanol and were transcardially perfused with PBS. Eyes were enucleated and fixed in 4% PFA in PBS for 18h at 4°C. Retinas were gently dissected from the eye cup and washed 3 times in PBS for 5 minutes, then washed 3 times in 0.3% Triton-X 100 in PBS for 5 minutes. Retinas were subsequently blocked in 10% Donkey Serum (Sigma-Aldrich, Cat#D9663) in 0.3% TritonX PBS for 24h at 4°C. Retinas were incubated in primary antibodies diluted in blocking buffer for 72h at 4°C and then washed 3× with PBS for 5 minutes. Retinas were then incubated with secondary antibodies in PBS for 24 hours at 4°C. Finally, retinas were washed 4 times with PBS for 5 minutes prior to whole mounting onto slides with fluorescent mounting medium (Polysciences, Cat#18606-20). Slides were allowed to dry prior to imaging. Primary antibodies utilized include: 1µg/mL Rabbit anti-RNA Binding Protein, MRNA Processing Factor (RBPMS) (GeneTex Cat#GTX118619). 0.5 μg/mL Donkey anti-Rabbit Alexa Fluor 568 (Invitrogen, Cat#A10042) was used as a secondary antibody.

Multicolor wide-field tiled images were taken on a Leica DMi8 microscope at 20× magnification across the entire retina. Retina images were stitched together in FIJI. All microscope settings were kept identical for each experiment. RBPMS^+^ RGC counts were determined by generating 4 separate 500µm x 500µm ROIs from the peripheral and central retina for each mouse. These ROIs were then subjected to automated thresholding using the moments algorithm (55), and watershed segmentation before counting all RBPMS^+^ objects above 100µm^2^. All analyses were performed with the Fiji distribution of ImageJ (NIH)(52, 53).

### Statistics

Analyses were performed using GraphPad Prism 10 software. Comparisons of percent of instances were analyzed using a Chi-square test. Data from experiments designed to test differences between two groups (e.g., one measurement across two ages within strain) were subjected to a Shapiro-Wilk test to test normality and an F test to compare variance. For normally distributed data with equal variance, a two-tailed independent samples *t* test was utilized. For normally distributed data with unequal variance, a Welch’s t test was used. For non-normally distributed data, a Mann-Whitney test was used. Data from experiments designed to test differences among more than two groups across one condition (e.g. one measurement across strains at one timepoint) were subjected to a Shapiro-Wilk test to test normality and a Brown-Forsythe test to compare variance. Normally distributed data with equal variance were analyzed using a one-way ANOVA followed by Tukey’s post-hoc test. Data from experiments designed to detect differences among multiple groups and across two (e.g. measurements across ages and sexes within strains) were analyzed using a two-way ANOVA followed by Holm-Sidak’s post-hoc test. *P* values <0.05 were considered statistically significant. Throughout the manuscript, results are reported as mean± standard error of the mean (SEM).

## Results

### Genetic background is a major source of transcriptomic and proteomic variation in the murine retina

Aging is an important risk factor for numerous vision-threatening diseases (5, 6), yet, little is known about the interaction of age, sex, and genetic background. To better understand how aging across diverse genetic contexts may alter retinal function, we profiled the retinal transcriptome and proteome of nine genetically diverse mouse strains at the ages of 4, 12 and ≥18M (Figure 1A). Several strains had natural attrition, reducing the number of aged mice of particular strains available for tissue collection and data analysis (see Supplemental Table 1 for full details). Our initial analysis of the resulting transcriptomic and proteomic data included sex, strain, and age as covariates. After filtering the multi-omics data, we analyzed 22,928 genes and 4,854 proteins across these groups. First, we visualized transcriptomic and proteomic data using principal component analyses (PCA). This revealed that strain was the largest contributor to variation in the transcriptomic data (Figure 1B) with more subtle patterns observed in the proteomic data (Figure 1C). The less pronounced strain signature in proteomic data may be due to the more limited numbers of proteins analyzed but may also be due in part to buffering at the protein level (56). As examples of the importance of strain variation in pathways critical for retinal health, genes associated with the GO term “Regulation of complement”, including *Cfh* (Figure 1D-E) or genes associated with the GO term “Detection of visible light”, including *Crb1,* displayed substantial strain-dependent variation (Figure 1F-G). Proteins within the “Metabolic process” GO term, including acetyl-CoA acetyltransferase 1 (ACAT1), also displayed strain-dependent expression patterns (Figure 1H-I). Thus, genetic context was a major source of variation in retinal molecular signatures.

**Figure 1.**
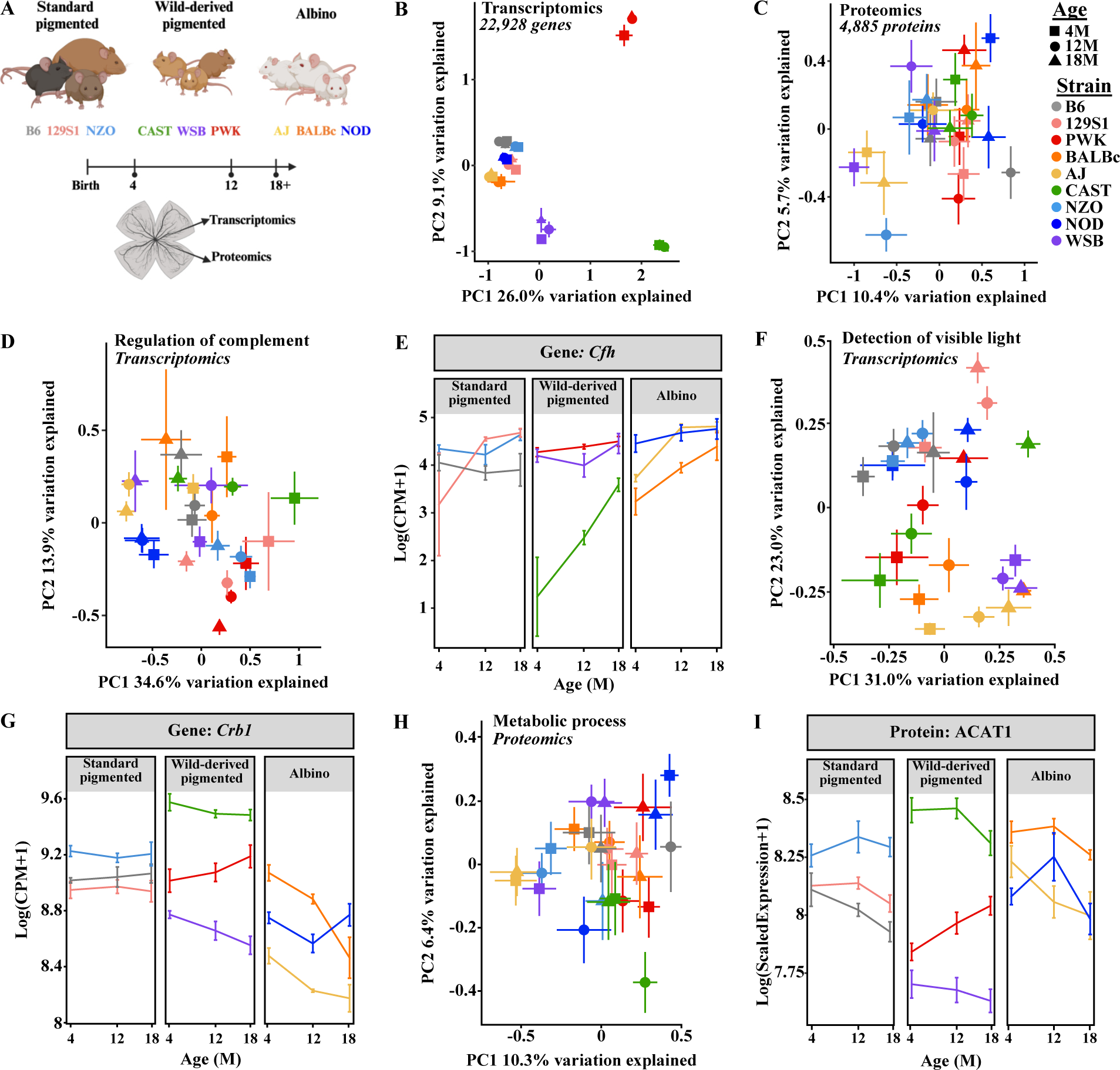
Multi-omics profiling of the aging retina across genetically diverse mice. **A.** Experimental design and schematic (created with BioRender.com). Principal component analysis (PCA) plots of **B.** transcriptomic data and **C.** proteomics data across all samples for PC1 and PC2. PCA plots and scatter plots visualizing changes in genes or proteins associated with the GO term: **D.** “Regulation of complement”, including **E**. *Cfh***, F.** “Detection of visible light”, including **G.** *Crb1*, and **H.** “Metabolic Process”, including **I.** ACAT1. Scatter plots in **E** and **G** illustrate changes in the log(counts per million mapped reads (CPM)+1) of *Cfh* and *Crb1*, and **I** illustrates the log(scaled peptide abundance) of ACAT1. In **B, D-G**: *N*=8 (4F,4M) NZO, 129S1, and B6 mice at each age; NOD mice: *N*=8 (4F,4M) at 4M*, N*=7 (3F,4M) at 12M, *N*=6 (2F,4M) at 18M; AJ mice: *N*=8 (4F,4M) at 4M and 12M, *N*=7 (4F,3M) at 18M; BALBc mice: *N*=7 (3F,4M) at 4M and 12M, *N*=4F at 18 M; PWK mice: *N*=8 (4F,4M) at 4M and 12M, *N*=3F at 18M; CAST mice: *N*=8 (4F,4M) at 4M and 12M, *N=7* (3F,4M) at 18M; WSB mice: *N=7* (3F,4M) at 4M, *N*=8 (4F,4M) at 12M and 18M. In **C, H, I**: *N*=8 (4F,4M) NZO, 129S1, and CAST mice at each age; NOD mice: *N*=7 (4F,3M) at 4M*, N*=8 (4F,4M) at 12M, *N*=6 (2F,4M) at 18M; AJ mice: *N*=8 (4F,4M) at 4M and 12M, *N*=7 (4F,3M) at 18M; BALBc mice: *N*=7 (3F,4M) at 4M, *N*=8 (4F,4M) at 12M, *N*=4F at 18 M; PWK mice: *N*=8 (4F,4M) at 4M and 12M, *N*=3F at 18M; B6 mice: *N*=8 (4F,4M) at 4M and 18M, *N=7* (3F,4M) at 12M; WSB mice: *N=7* (3F,4M) at 4M, *N*=7 (4F,3M) at 12M, *N*=8 (4F,4M) at 18M. In **B-I**: error bars represent SEM.

### The aging retinal transcriptome and proteome are influenced by genetic context

To define an overall retinal aging signature regardless of genetic context, we tested for differential gene and protein expression using age as the predictive variable across genetic backgrounds. We found that 4929 genes and 271 proteins were differentially expressed with aging across genetically diverse mice (Supplemental Figure 1A-B). Of these 271 differentially expressed proteins (DEPs), 131 were also significantly modulated at the gene level (Supplemental Figure 1C). Surprisingly, sex did not have a significant impact on the aging signatures across strains. Enrichment analyses revealed significant alteration of GO terms involving immune processes, photoreceptors, metabolism, and chromatin organization at both the gene and protein level (Supplemental Figure 1D-E). Differentially expressed photoreceptor genes and proteins decreased with age, while the differentially expressed genes (DEGs) and DEPs associated with immune processes largely increased with age (Supplemental Table 2).

Next, we sought to examine how this aging signature was modified by genetic context. PCA visualization of the aging signature genes and proteins suggested differences in gene and protein expression among strains of mice (Figure 2A-B). As examples, genes associated with the GO term “Regulation of histone methylation”, including *Mthfr* (Figure 2C-D), genes associated with the GO term “Antigen processing and peptide presentation”, including *B2m* (Figure 2E-F), and genes associated with the GO term “Retina homeostasis”, including *Rho* (Figure 2G-H) all displayed substantial strain- andage-dependent variation. The observed differences in the aging signature suggested these strains may display differential susceptibility to aging-associated retinal diseases.

**Figure 2.**
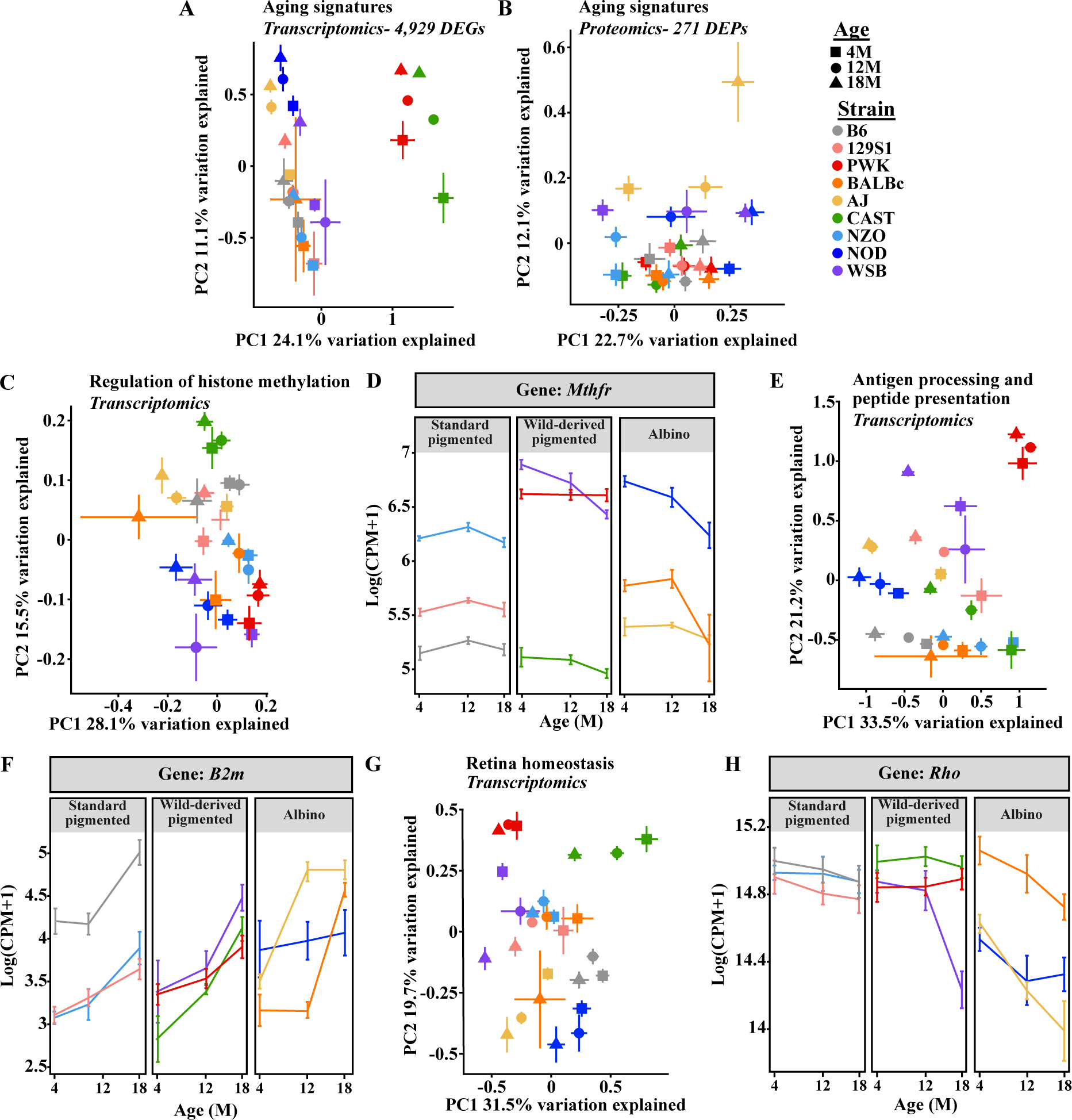
Genetic context influences expression of proteins and genes associated with aging. Principal component analysis (PCA) plots of the common aging signature across all strains and sexes identified in **A.** transcriptomic data and **B.** proteomics data. PCA plots and scatter plots visualizing changes in aging signature enriched genes or proteins associated with the GO term: **C.** “Regulation of histone methylation”, including **D**. *Mthfr*, **E.** “Antigen processing and peptide presentation”, including **F**. *B2m*, and **G.** “Retina homeostasis”, including **H**. *Rho*. For **D,F,H**: Scatter plots illustrating changes in the log(counts per million mapped reads (CPM)+1). In **A,C-H**: *N*=8 (4F,4M) NZO, 129S1, and B6 mice at each age; NOD mice: *N*=8 (4F,4M) at 4M*, N*=7 (3F,4M) at 12M, *N*=6 (2F,4M) at 18M; AJ mice: *N*=8 (4F,4M) at 4M and 12M, *N*=7 (4F,3M) at 18M; BALBc mice: *N*=7 (3F,4M) at 4M and 12M, *N*=4F at 18 M; PWK mice: *N*=8 (4F,4M) at 4M and 12M, *N*=3F at 18M; CAST mice: *N*=8 (4F,4M) at 4M and 12M, *N=7* (3F,4M) at 18M; WSB mice: *N=7* (3F,4M) at 4M, *N*=8 (4F,4M) at 12M and 18M. In **B**: *N*=8 (4F,4M) NZO, 129S1, and CAST mice at each age; NOD mice: *N*=7 (4F,3M) at 4M*, N*=8 (4F,4M) at 12M, *N*=6 (2F,4M) at 18M; AJ mice: *N*=8 (4F,4M) at 4M and 12M, *N*=7 (4F,3M) at 18M; BALBc mice: *N*=7 (3F,4M) at 4M, *N*=8 (4F,4M) at 12M, *N*=4F at 18 M; PWK mice: *N*=8 (4F,4M) at 4M and 12M, *N*=3F at 18M; B6 mice: *N*=8 (4F,4M) at 4M and 18M, *N=7* (3F,4M) at 12M; WSB mice: *N=7* (3F,4M) at 4M, *N*=7 (4F,3M) at 12M, *N*=8 (4F,4M) at 18M. In **A-H**: error bars represent SEM.

Common retinal neurodegenerative diseases involve RGC and/or photoreceptor loss (6, 12, 57). Single nucleus RNA-sequencing datasets have revealed several marker genes for specific retinal neurons (18). Therefore, we probed for changes in these neuronal populations by visualizing marker gene differences. We first profiled changes in photoreceptor-specific gene expression across strains using PCA (Figure 3A). We found that WSB, NOD, and AJ mice showed age-associated shifts away from the other strains (Figure 3A). In fact, these three strains exhibited reduced photoreceptor gene expression relative to the other strains at 18M (Figure 3B). In accordance with our transcriptomics data, NOD and AJ mice are known to develop age-associated loss of photoreceptors (58, 59). In our analysis, WSB mice exhibited age-dependent differences in photoreceptor marker gene expression (Figure 3C). WSB mice do not carry mutations in known retinal degeneration genes (60), and to date, there are no published reports of WSB photoreceptor loss. Thus, WSB mice may serve as a novel model of photoreceptor degeneration.

**Figure 3.**
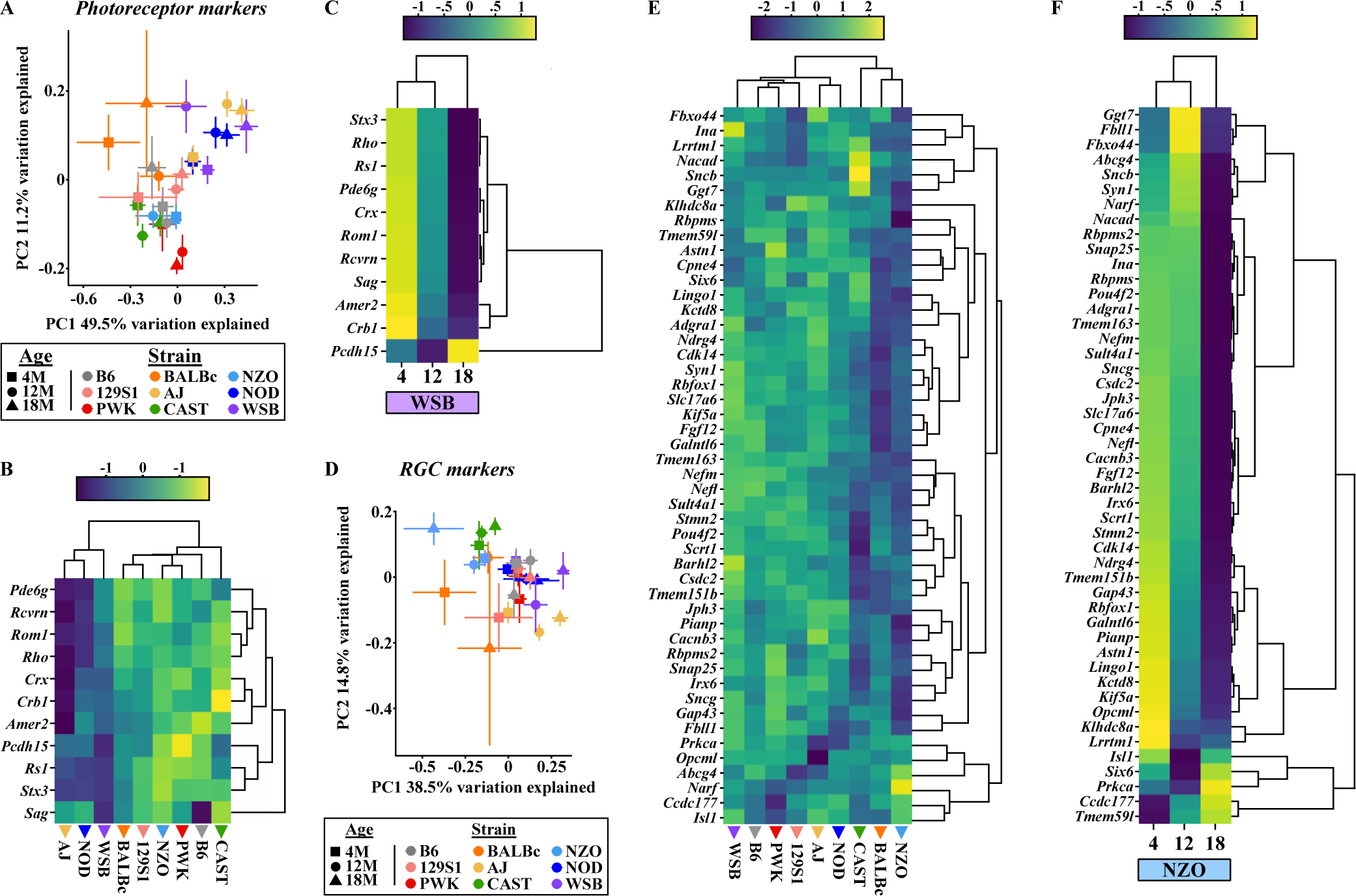
Photoreceptor and RGC markers change in strain-specific manners across aging. **A**. Principal component analysis (PCA) plot of the expression of unique marker genes associated with photoreceptors across all strains. Heatmaps of marker genes associated with photoreceptors across all strains at 18M **B.** and across aging in **C.** WSB mice. **D.** PCA plot of the expression of unique marker genes associated with RGCs across all strains. Heatmaps of marker genes associated with RGCs across **E.** all strains at 18M and across aging in **F.** NZO mice. In **B,C,E, F**: average expression for the entire group is shown. Scaled and centered expression of the library-size and log normalized average expression value for each gene in each group. In **A,B,D, E**: *N*=8 (4F,4M) NZO, 129S1, and B6 mice at each age; NOD mice: *N*=8 (4F,4M) at 4M*, N*=7 (3F,4M) at 12M, *N*=6 (2F,4M) at 18M; AJ mice: *N*=8 (4F,4M) at 4M and 12M, *N*=7 (4F,3M) at 18M; BALBc mice: *N*=7 (3F,4M) at 4M and 12M, *N*=4F at 18 M; PWK mice: *N*=8 (4F,4M) at 4M and 12M, *N*=3F at 18M; CAST mice: *N*=8 (4F,4M) at 4M and 12M, *N=7* (3F,4M) at 18M; WSB mice: *N=7* (3F,4M) at 4M, *N*=8 (4F,4M) at 12M and 18M. In **C**: *N=7* (3F,4M) WSB mice at 4M, *N*=8 (4F,4M) WSB mice at 12M and 18M. In **F**: *N*=8 (4F,4M) NZO at each age. In **A,D**: error bars represent SEM.

In addition to photoreceptors, we profiled changes in RGC marker genes across strains. NZO, CAST, and BALBc mice exhibited a slight leftward shift in the PCA plot for RGC genes (Figure 3D), and heatmap visualization of RGC marker genes revealed low expression of these genes in aged NZO, CAST and BALBc mice relative to other strains (Figure 3E). A previous report suggested CAST mice develop significantly fewer RGCs (and thus have smaller optic nerves) than other strains, which may account for the reduced expression of RGC genes in this strain (61). BALBc mice have been shown to develop mild age-related retinal dysfunction (61). However, there have been no previous reports of retinal phenotypes in NZO mice. To better understand how aging influences RGC gene expression in NZO mice, we visualized gene expression changes across age in NZO mice and found striking reductions in these genes (Figure 3F). These data suggest NZO mice may be especially susceptible to RGC loss with age.

Collectively, our multi-omics analysis suggested genetic context significantly contributes to retinal aging signatures. We created a user-friendly web application to increase data accessibility and foster hypothesis development and refinement, which can be accessed at: https://thejacksonlaboratory.shinyapps.io/Howell_AgingRetinaOmics/.

### NZO and WSB mice exhibit disease-relevant retinal fundus abnormalities with age

Our analyses suggested shared changes to photoreceptor markers across all nine strains, along with previously unreported age-dependent changes to retinal neurons in NZO and WSB mice. We therefore sought to verify these predictions using a battery of clinically-relevant *in vivo* ophthalmological exams at the ages of 4, 8, 12 and/or ≥18M in NZO, WSB, and B6 mice of both sexes (Figure 4A). Gross changes to the anterior segment of NZO, WSB, and B6 eyes were not detected using slit lamp imaging (Figure 4B). In contrast, fundus imaging revealed age-associated increases of fundus abnormalities in WSB, NZO, and B6 mice (Figure 4C-D). Fundus abnormalities in B6 mice included abnormal retinal spots (Figure 4C-D). In NZO mice, these abnormalities included spots resembling cotton wool spots and exudates, which are commonly observed in human DR patients and other retinal vascular diseases (62, 63). The presence of these phenotypes peaked by 12M, however, what appeared to be an epiretinal membrane was uniquely observed in aged NZO eyes (7 of 38 NZO 18M eyes; Supplemental Figure 2). WSB mice exhibited fundus abnormalities as early as 4M, which became increasingly severe with age (Figure 4C-D). These abnormalities included potential pigment disruption, retinal spots, and waxy optic disc pallor. Altogether, these data indicate age-related changes to DR-relevant fundus phenotypes in NZO, and retinal degeneration-relevant phenotypes in WSB mice.

**Figure 4.**
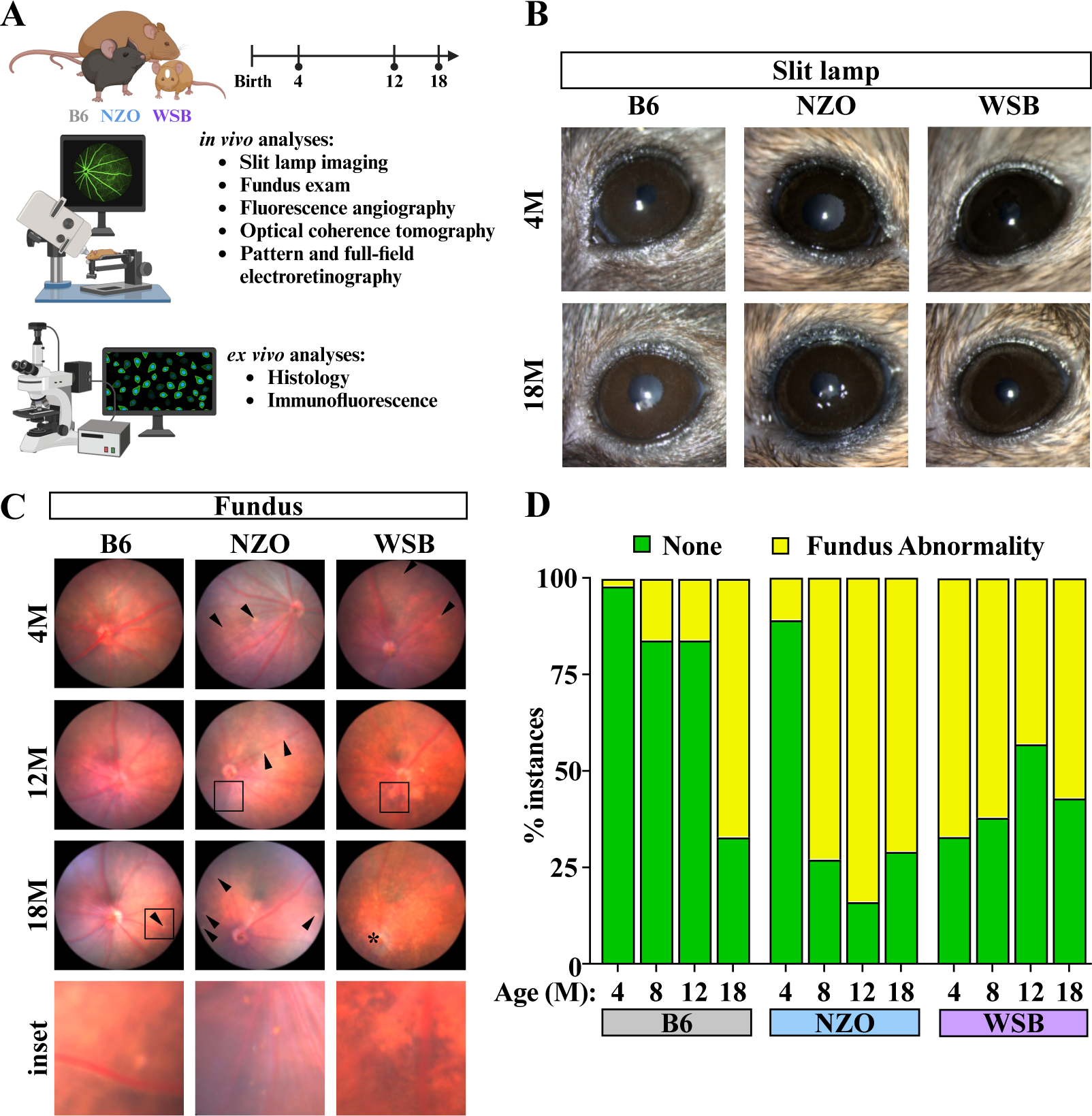
Validating Omics analyses using in vivo characterization. **A**. Experimental design schematic (created with BioRender.com). **B**. Representative slit lamp images of 4 and 18M B6, NZO and WSB mice. **C.** Representative fundus images of B6, NZO and WSB retinas at the ages of 4, 12, and 18M. Arrowheads indicate fundus spots. * indicates waxy optic disc pallor. Insets highlight different fundus abnormalities: B6 – fundus spot, NZO – exudates, WSB – pigmentary change. **D.** Quantification of the proportion of eyes affected by fundus abnormalities at 4, 8, 12 and 18M. In **D**: chi-square tests were used to test differences between ages within each strain: B6 – *p*<0.0001; NZO – *p*<0.0001; WSB – *p=*0.0044. For B6: *N*=156 eyes at 4M, 64 eyes at 8M, 63 eyes at 12M, and 24 eyes at 18M. For NZO: *N*=160 eyes at 4M, 70 eyes at 8M, 64 eyes at 12M, and 31 eyes at 18M. For WSB: *N*=36 eyes at 4M, 42 eyes at 8M, 60 eyes at 12M, and 46 eyes at 18M.

### OCT profiling unveils strain-specific patterns of retinal thinning

To investigate whether fundus abnormalities corresponded with gross retinal degeneration, OCT was employed to measure retinal layer thicknesses across age in each strain (Figure 5A). As total retinal thickness differed by strain in young animals (Figure 5A-B), we determined the effect of age within each strain separately. Age reduced total retinal thickness across all three strains (Figure 5B). There was a subtle but significant interaction effect between sex and age in NZO mice (Figure 5B). The nerve fiber layer (NFL), ganglion cell layer (GCL), and inner plexiform layer (IPL) are largely composed of RGC axons, cell bodies, and dendritic processes/synapses, respectively. These layers were difficult to discern via OCT due to technical limitations, therefore, the NFL, GCL, and IPL were measured as one complex (NFL/IPL/GCL). For the NFL/IPL/GCL complex, there was a significant age effect in WSB mice (Figure 5C) and an age-by-sex effect in NZO mice (Figure 5C). We found significant age-associated reductions in the inner nuclear layer (INL) and outer nuclear layer (ONL) in NZO and B6 mice (Figure 5D-E). WSB mice exhibited age- and sex-by-age driven reductions in the INL and ONL, respectively (Figure 5D-E). In brief, 1) B6 mice lost 6% total retinal thickness by 18M, which was due to a 19% loss of INL thickness and 12% loss of ONL thickness; 2) NZO mice also lost 6% total retinal thickness by 18M, which was due to a 21% loss of INL thickness and 17% loss of ONL thickness; 3) WSB mice lost 22% total retinal thickness by 18M, which was due to a 19% loss of INL thickness and 49% loss of ONL thickness. As sex was associated with such subtle effects, and no significant effects in the aging signatures within the multi-omics analyses were observed, we combined sexes for future analyses.

**Figure 5.**
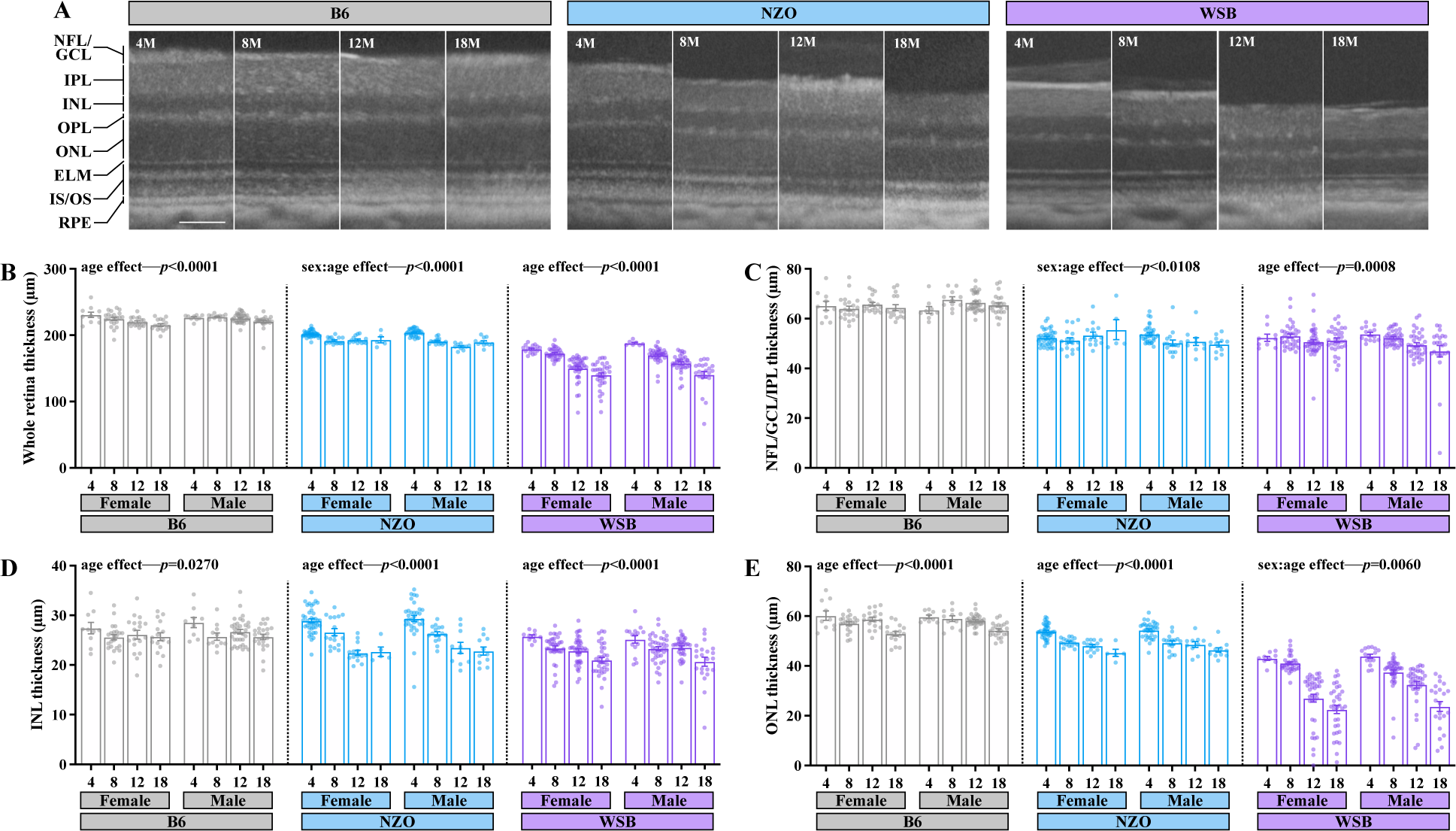
Optical Coherence Tomography (OCT) reveals strain specific aging effects on retinal thinning. **A**. Representative OCT images at 4, 8, 12 and 18M of age in B6, NZO and WSB mice. NFL: nerve fiber layer, GCL: ganglion cell layer, IPL: inner plexiform layer, INL: inner nuclear layer, OPL: outer plexiform layer, ONL: outer nuclear layer, ELM: external limiting membrane, IS/OS: inner/outer segment, RPE: retinal pigment epithelium. Scale bar, 100µm. Quantification of: **B.** the total retina thickness (distance from top of NFL/GCL to bottom of the RPE) across strains and sexes and ages, **C.** NFL/GCL/IPL complex thickness, **D.** INL thickness, and **E.** ONL thickness. In **B-E**: due to prominent differences in retinal organization by strain, two-way ANOVAs within each strain were utilized to evaluate the impact of aging and sex. For B6: *N*=19 (10F,9M) eyes at 4M, 33 (22F,11M) at 8M, 51 (18F,33M) at 12M, 39 (16F,23M) at 18M; for WSB: *N*=24 (17F,7M) eyes at 4M, 72 (34F,38M) at 8M, 77 (40F,36M) at 12M, 55 (33F,22M) at 18M; for NZO: *N*=73 (40F,33M) eyes at 4M, 33 (17F,16M) at 8M, 24 (14F,10M) at 12M, 16 (5F,11M) at 18M. Error bars represent SEM.

Altogether, these data suggest that aging mice exhibit neural retinal thinning regardless of genetic context but that the specific age- and sex-affected disruptions are strongly influenced by genetics. Moreover, these perturbations were strongest in WSB mice, which may disproportionally lose photoreceptors within the ONL.

### WSB mice exhibit abnormal oscillatory potentials and exacerbated aging-associated reductions in photoreceptor function

Given the extensive thinning of the ONL in WSB mice, we next sought to evaluate photoreceptor function with scotopic and photopic ERGs (dark-adapted and light-adapted, respectively) at 4 and 18M (Figure 6). As was predicted by our omics analyses, age reduced these amplitudes across strains (Figure 6A-I). At a luminance that elicited maximal ERG amplitudes (10cd.s/m^2^), WSB mice exhibited a 52% decline in scotopic a-wave amplitudes, while B6 and NZO mice had 15% and 26% reductions, respectively (Figure 6B-C). These data suggest drastic rod dysfunction in aged WSB mice. Scotopic b-waves were reduced by 34% in WSB mice, 17% in B6 mice, and 27% in NZO mice (Figure 6D-E). Both NZO and WSB mice had an age-associated decrease in scotopic oscillatory potentials (OPs; 24% and 37%, respectively), which was not observed in B6 mice (Figure 6F-G). Strikingly, WSB mice appeared to exhibit nearly absent OPs even at young ages. In contrast, photopic b-wave amplitudes were similar across strains in young mice, and WSB and NZO mice exhibited mild age-associated reductions in photopic b-wave amplitudes (20% and 25%, respectively; Figure 6H-I). These data suggest relatively conserved cone function across strains with age.

**Figure 6.**
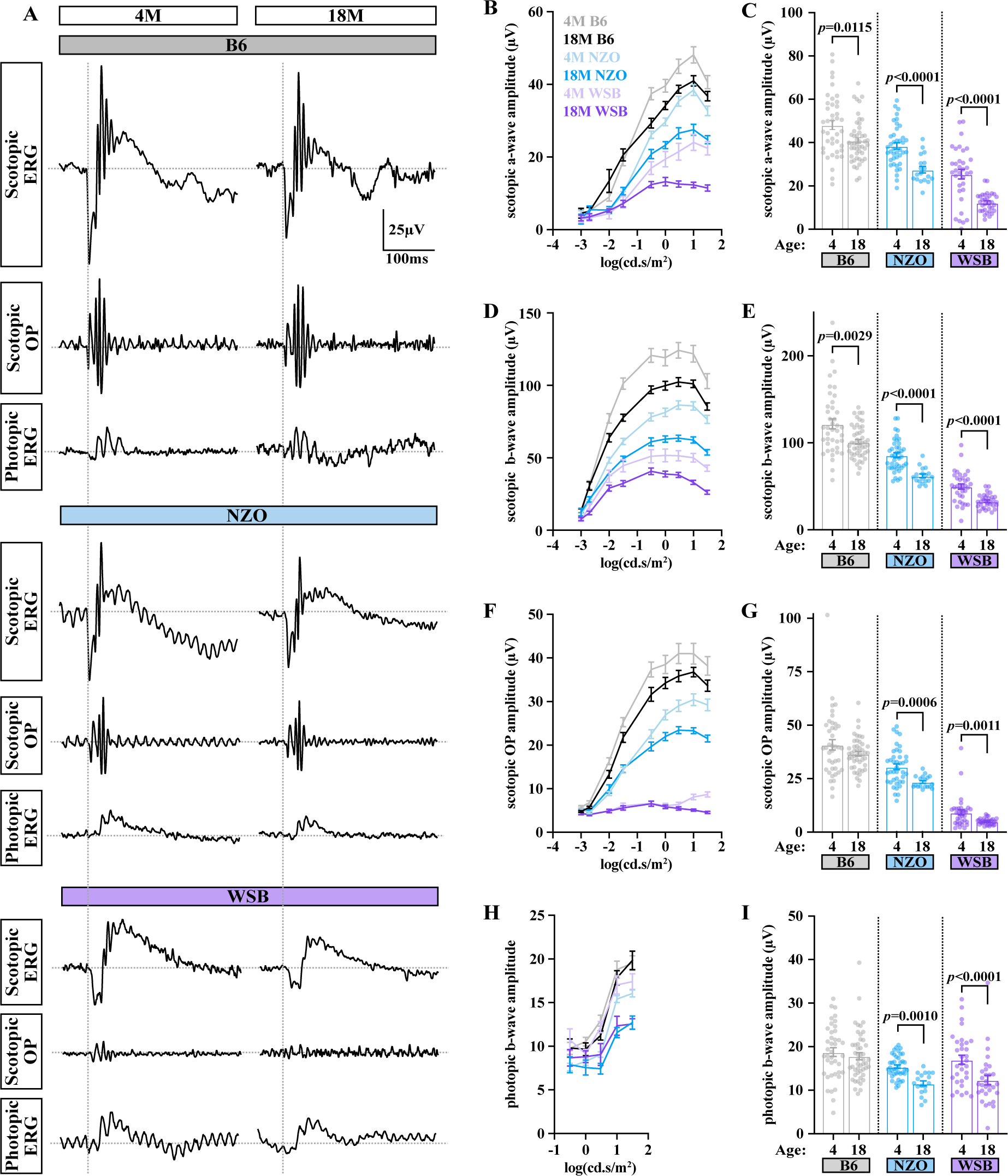
WSB mice exhibit enhanced aging-associated photoreceptor functional decline. **A**. Representative scotopic and photopic ERG and OP curves at 10 cd.s/m^2^ luminance in B6, WSB and NZO mice. Quantification of scotopic **B.** a-wave amplitude (µV), **D.** b-wave amplitude (µV), and **F.** OP amplitude (µV) across luminance intensities for 4M and 18M B6, NZO and WSB mice. Amplitudes at 10 cd.s/m^2^ luminance for 4 and 18M B6, NZO, and WSB mice for the scotopic **C.** a-waves, **E.** b-waves, **G.** OPs. **H.** Quantification of photopic b-wave amplitude (µV) across luminance intensities for 4M and 18M B6, NZO and WSB mice. **I**. Quantification of photopic b-wave amplitudes (µV) at 10 cd.s/m^2^ luminance for 4 and 18M B6, NZO, and WSB mice. Error bars represent SEM. In **A-G**: for B6: *N*=40eyes at 4M, 48 at 18M; for WSB: *N*=37 eyes at 4M, 32 at 18M; for NZO: *N*=40 eyes at 4M, 18 at 18M. In **H-I**: for B6: *N*=40eyes at 4M, 46 at 18M; for WSB: *N*=32 eyes at 4M, 30 at 18M; for NZO: *N*=40 eyes at 4M, 18 at 18M. All comparisons were made within strains due to baseline differences. In **C**: for B6: Mann-Whitney U test; for NZO: Student’s t-test; for WSB: Welch’s t-test. In **E**: for B6: Mann-Whitney U test; for NZO and WSB: Welch’s t-test. In **G**: for all three strains: Mann-Whitney U test. In **I**: for B6 and NZO: Mann-Whitney U test; for WSB, Student’s t-test.

Given drastic ONL thinning and rod photoreceptor dysfunction in WSB mice, we next sought to quantify photoreceptor cell loss more precisely, as OCT images may not capture changes outside of the central retina due to technical limitations. Thus, next employed routine hematoxylin and eosin (H&E) on retinal sections at 4, 12, and 18M in B6, NZO, and WSB mice (Figure 7A). In accordance with OCT measurements, the number of ONL cells decreased with age across strains (Figure 7B). B6 retinas had a 12-13% decrease in ONL cells in the center and middle retina by 12M, which declined further by 15-29% across the retina by 18M. NZO retinas had a 19-23% loss of ONL cells by 12M, which did not decline further by 18M. WSB retinas had a 14-22% loss of ONL cells by 12M, but further lost 37-59% of ONL cells by 18M. This loss was most drastic in the center retina. In fact, we observed nearly complete loss of ONL cells in the central retina of 18M WSB animals alongside instances of abnormal retinal structures (Supplemental Figure 3).

**Figure 7.**
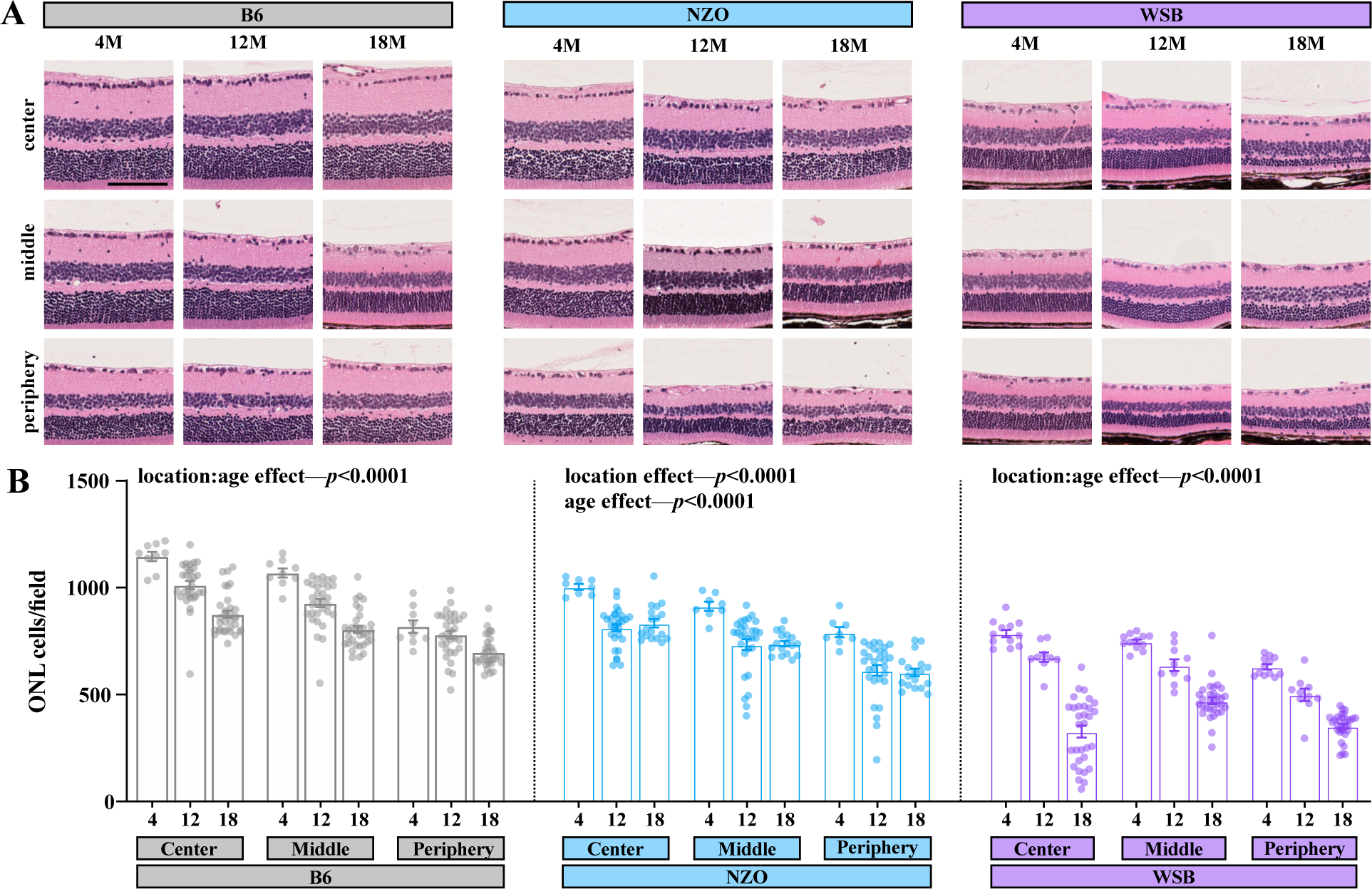
Aging is associated with outer nuclear layer cell loss, which is exacerbated in WSB animals. **A.** Representative hematoxylin and eosin (H&E) staining of retinas at three anatomical locations relative to the optic nerve head across B6, NZO, and WSB mice at 4,12, and 18M. **B.** Automated cell counts within the outer nuclear layer (ONL). In **B**: mixed effects analyses with repeated measures were used to assess regional and aging effects within strains. Error bars represent SEM. In **B**: for B6: *N*=9 eyes at 4M, 31 at 12M, 32 at 18M; for WSB: *N*=12 eyes at 4M, 8-10 at 12M, 31 at 18M; for NZO: *N*=8 eyes at 4M, 29 at 12M, 17-18 at 18M.

These assays indicate that aging reduces rod photoreceptor function across all three mouse strains, and WSB mice exhibit early retinal dysfunction with severe rod dysfunction and drastic loss of central ONL cells with age.

To begin to understand the mechanisms driving the early and progressive photoreceptor dysfunction in WSB mice, we compared proteome differences between 4M WSB with other pigmented strains. This approach revealed that 258 proteins were altered in WSB mice relative to the other strains and were associated in pathways involving essential mitochondrial metabolism, including “Generation of precursor metabolites and energy”, “Purine ribonucleotide metabolic process”, and reductions to proteins associated with the BBSome (Supplemental Figure 4A-B, Supplemental Table 3). The BBSome is critical for photoreceptor outer segment function, and loss of BBSome proteins have been shown to cause photoreceptor degeneration and dysfunction (64-66). Our data show reduced expression of WSB BBSome proteins relative to other strains even at a young age. Furthermore, Retinol Dehydrogenase 8 (the major outer segment enzyme responsible for converting all-*trans*-retinal to all-*trans*-retinol and preventing accumulation of toxic by-products) was significantly lower in WSB retinas (67). Thus, early WSB photoreceptors dysfunction may be due in part to cell-intrinsic deficits in outer segment functions.

### NZO mice display profound loss of RGC function with age and exhibit retinal microvascular dysfunction

Our transcriptomic data suggested NZO mice lose RGCs with age (Figure 3D-F). Thus, we utilized PERG to probe for gross changes in RGC potentials (68). At 4M, NZO mice exhibited blunted PERG amplitudes (Figure 8A-B), which declined by 41% to near-noise levels by 18M—indicating profound RGC dysfunction. Interestingly, B6 mice displayed a 28% reduction in PERG amplitudes by 18M, while WSB mice were resilient to age-related PERG decline. To determine whether RGCs are lost (versus less functional), we performed whole-mount immunohistochemistry for RNA Binding Protein, MRNA Processing Factor (RBPMS)+ cells to quantify this neuronal population across the retina. We found that 12 and 18M NZO mice had a ∼35-56% decrease in surviving RGCs relative to 4M NZO animals, compared to 5-13% and 25-26% for B6 and WSB, respectively (Figure 8C-D). Collectively, these data indicate NZO mice display RGC dysfunction and loss.

**Figure 8.**
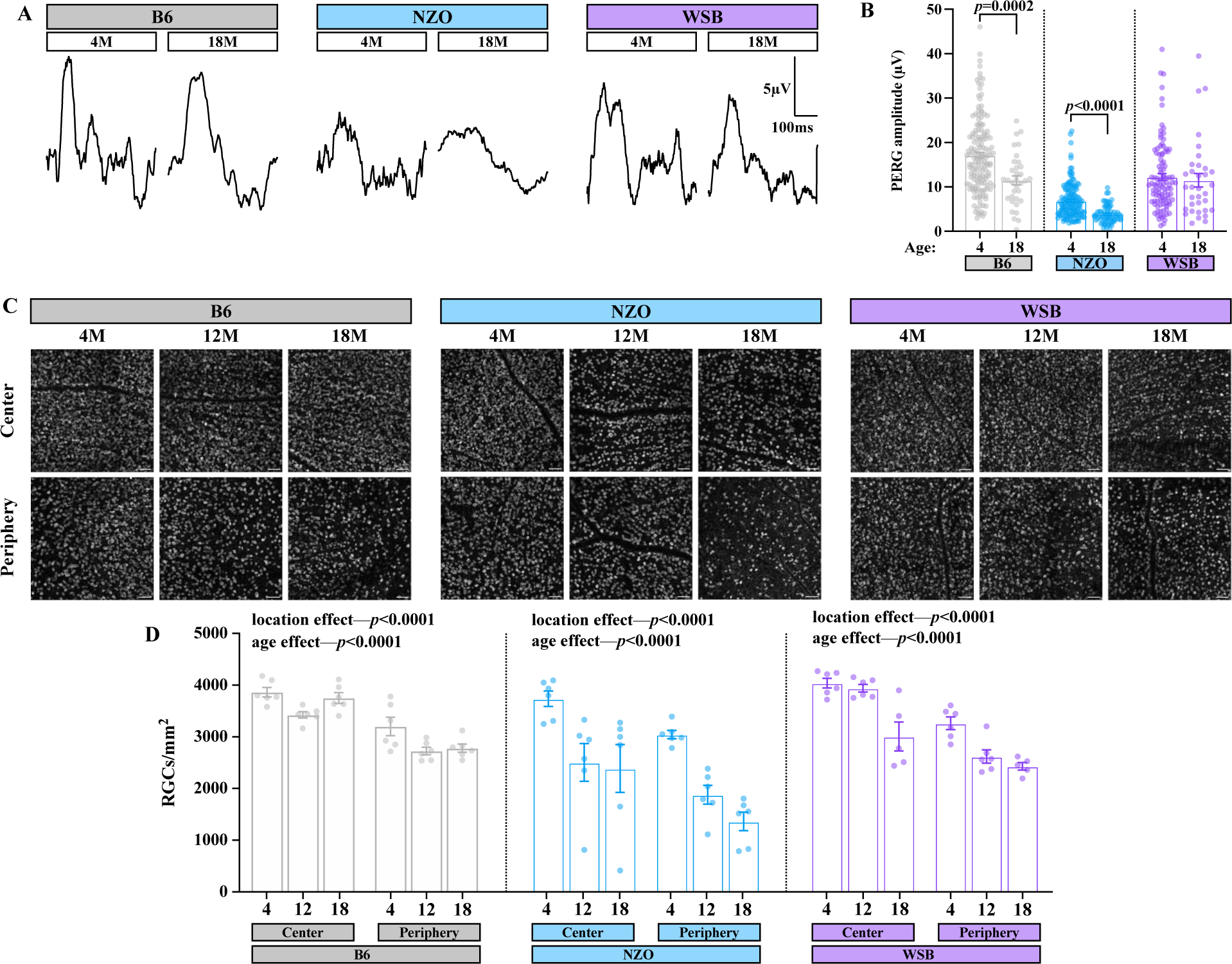
NZO mice exhibit reduced gross RGC potentials and RGC loss with age. **A**. Representative PERG response curves from B6, WSB and NZO mice at 4 and 18M. **B.** Amplitude of PERG response (µv) in 4M and 18M B6, WSB and NZO mice. **C.** Representative images of RBMPS immunohistochemistry in B6, NZO, and WSB mice at 4, 12, and 18M. Scale bar:50µm. **D.** Quantification of automated RBPMS+ retinal ganglion cells (RGCs) counts per mm^2^ in the central retina and peripheral retina. In **A-B**: for B6: *N*=152 eyes at 4M, 34 at 18M; for WSB: *N*=101 eyes at 4M, 34 at 18M; for NZO: *N*=142 eyes at 4M, 66 at 18M. In **C-D:** for B6 and NZO: *N*=6 eyes at 4,12 and 18M; for WSB: *N*=6 eyes at 4 and 12M, 5 at 18M. In **B**: Mann-Whitney U tests were used to evaluate differences between ages within strains. In **D**: two-way ANOVAs were used to assess effects of region and age within strains. Error bars represent SEM.

In addition to apparent RGC loss, aged NZO mice have been shown to exhibit most features of metabolic syndrome (69-71). It is well established that retinal microvascular dysfunction coincides with metabolic stress— including diabetes and hypertension (72, 73). Therefore, we next evaluated retinal vascular changes using fluorescein angiography *in vivo* in B6, NZO, and WSB mice. We found differences in the mean intensity of fluorescein dye across strains (Figure 9A-C). Interestingly, age significantly reduced the mean intensity across all three strains, with a dramatic effect observed in NZO mice (Figure 9A-C). B6 and WSB mice had a 70% and 60% reduction in mean fluorescence intensity of fluorescein, respectively, while NZO mice had an 86% reduction in intensity. We also observed several instances of microaneurysms and hemorrhages in NZO animals by fundus and angiography (Supplemental Figure 5A-E). Furthermore, we detected a few instances of Prussian Blue+ deposits, indicating subtle intraretinal vascular leakage in aging NZO mice (Supplemental Figure 5F). Collectively, these data suggested that NZO animals exhibit reduced retinal perfusion with age or reduced capillary density. To further investigate retinal vascular deficits, we utilized a FITC-dextran permeability assay to probe the integrity of the blood-retina barrier and found that NZO exhibited significantly higher instances of 70kDa FITC-dextran leakage. We noted increased areas of nonperfusion in NZO relative to either B6 or WSB mice at 12M of age (Figure 9D-E). To determine if these areas of nonperfusion were associated with capillary loss, we isolated and stained retinal vascular networks from 12M animals. Wefound there was significantly more acellular retinal capillaries in 12M NZO retinas (41 acellular capillaries/mm^2^) compared to B6 (21/mm^2^) and WSB (18/mm^2^) retinas(Figure 9F-G). Altogether, these data suggest NZO mice exhibit DR-relevant microvascular dysfunction and capillary loss by 12M of age, which may influence RGC loss and progression of neurodegeneration.

**Figure 9.**
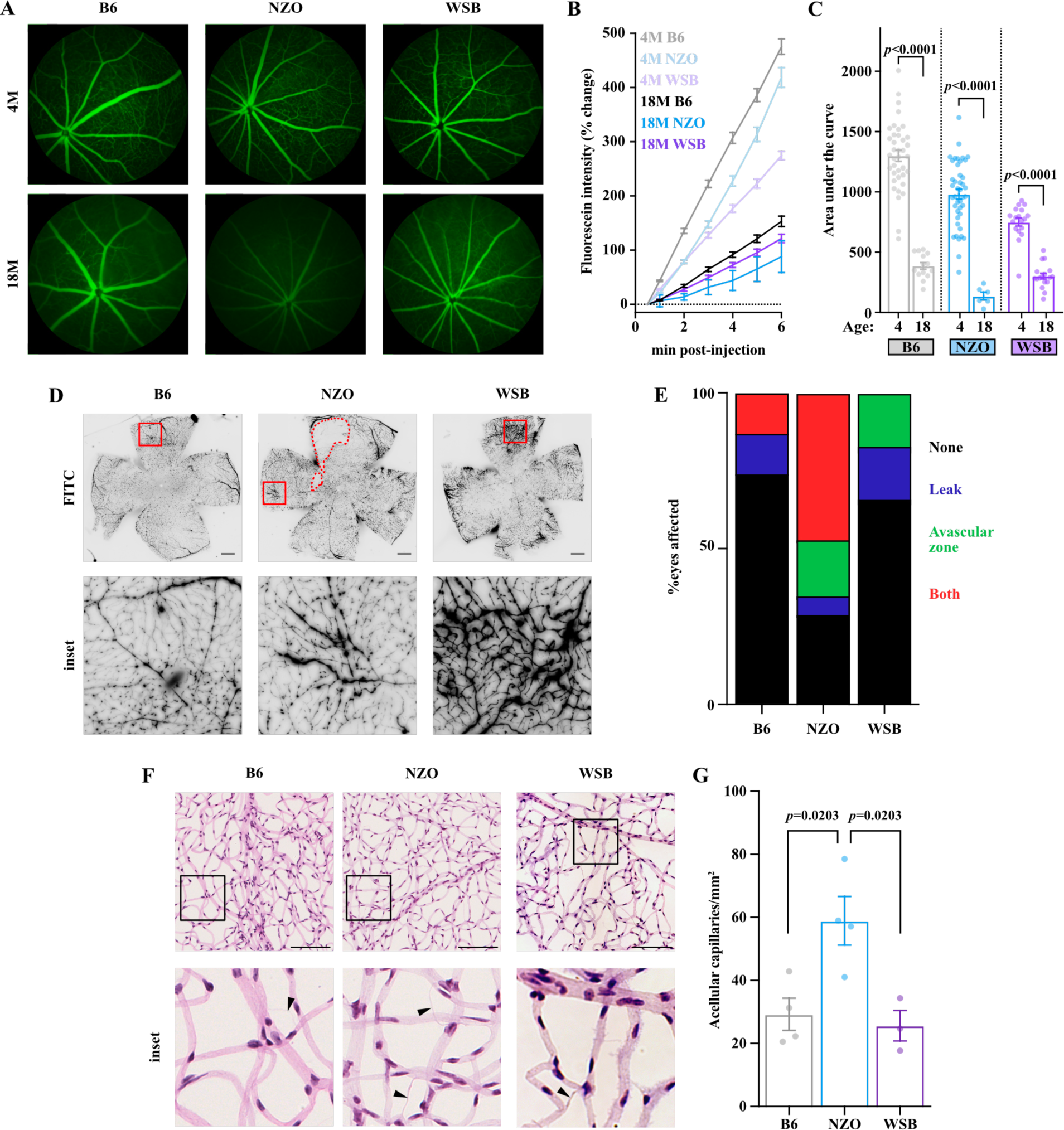
NZO mice exhibit microvascular dysfunction and capillary loss. **A.** Representative images of fluorescein angiography at 6minutes post administration in B6, NZO, and WSB mice at 4 and 18M. **B.** Quantification of change in fluorescent intensity of fluorescein from initial measurement across groups. **C.** Quantification of the area under the curve from **B** for B6, NZO, and WSB mice at 4 and 18M. **D.** Representative images of 70kDa FITC-Dextran leakage assay within 12M B6, NZO and WSB mice. Insets highlight notable findings. Scale bar, 100µm. **E.** Quantification of presence of vascular abnormalities in B6, NZO, and WSB mice. Chi-square test: *p*<0.0001. **F.** Representative images of retinal vascular networks stained with H&E in 12M B6, NZO and WSB mice. Insets highlight areas with acellular capillaries (arrowheads). Scale bar: 100µm. **G.** Quantification of acellular capillaries per mm^2^ in 12M B6, NZO and WSB mice. In **A-C**: for B6: *N*=38 eyes at 4M, 32 eyes at 8M, 30 at 12M, 14 at 18M; for WSB: *N*=18 eyes at 4M, 22 at 8M, 19 at 12M, 17 at 18M; for NZO: *N*=42 eyes at 4M, 35 at 8M, 30 at 12M, 6 at 18M. For **D-E**: *N*=8 B6, 17 NZO and 6 WSB eyes. For **F-G**: *N*=4 B6, 4 NZO and 3 WSB eyes. In **C**: for B6 and NZO: Welch’s t-test; for WSB: Mann-Whitney U test. In **G**: one-way ANOVA (*p*=0.0101) with Tukey’s multiple comparison test. Error bars represent SEM.

To identify potential mechanisms driving increased susceptibility to RGC loss in NZO mice, we compared proteome differences between NZO mice and the other pigmented strains at 4M. This approach revealed that 97 proteins were altered in NZO mice relative to the other strains, which were associated with antioxidant pathways such as “Glutathione metabolic process” including the protein Glutathione Synthetase (GSS), “Cellular detoxification” including the enzyme Glyoxalase 1 (GLO1), and changes to proteins associated with the NADH oxidoreductase complex (Supplemental Figure 6A-B, Supplemental Table 4). These data indicate NZO retinas may be particularly susceptible to oxidative stress associated with metabolic dysfunction due to reduced protective capacities.

## Discussion

Vision-threatening retinal degenerative disorders are becoming increasingly prominent within the aging population. Age is a critical risk factor in the development of retinal neurodegeneration. Studies in both humans and mice have demonstrated that genetics influences susceptibility to age-related retinal degeneration. However, few studies have investigated molecular changes occurring in the retina with age, or how these changes may be modified by genetic context. Elucidating these molecular changes will be critical in improving our understanding of retinal aging and disease, potentially informing novel models and therapeutic targets. Furthermore, these data can identify mechanisms that are critical for neurodegenerative diseases that impact the brain and are an essential foundation to determine if retinal changes can be used as a biomarker for brain neurodegeneration. In the present study, we investigate retinal transcriptomic and proteomic changes with age across nine genetically diverse mouse strains. This diverse population, with the inclusion of both sexes across young (4M), middle-aged (12M), and aged (18M+) mice, is intended to improve the relevance of these data to the heterogeneous human population.

Our multi-omic investigation revealed common aging signatures across strains, which included reduced photoreceptor and visual processing functions concomitant with activation of immune signaling pathways at both the gene and protein level. In concordance with this, we found subtle but significant loss of ONL cells and a decline in scotopic ERG a- andb-waves across strains measured, including the widely utilized B6 mouse. These data indicate subtle age-related loss of rod photoreceptors and INL cells, respectively. Our results align with the reduced ERG function and photoreceptor loss observed in human patients with age (74). These data suggest that even in the absence of frank neurodegeneration, there is increasing neural dysfunction with age in the murine retina. It is unclear whether the activation of immune response pathways is in response to neural dysfunction, or if immune activation is directly responsible for the observed dysfunction. Future studies will be required to probe these possibilities.

Strain was the strongest contributor to variation within the transcriptomic and proteomic data. In our analyses, we highlight that genetic context dictated the expression patterns of genes and proteins associated with a conserved aging signature. For example, genes associated with conserved aging-induced “Antigen processing and peptide presentation” pathway displayed substantial strain-dependent differences, with PWK mice exhibiting less age-dependent changes in these genes. The identified common aging signature across mice suggested altered visual function with age. However, some strains exhibited greater changes in these pathways, including those associated with photoreceptors. While previous work has linked aging to dysregulation of retinal neurons, overt loss of photoreceptors and RGCs is intimately associated with many human diseases (12, 68, 75). To infer changes in photoreceptor and RGC populations with age, we visualized the expression of cell-type specific marker genes across ages. Unfortunately, many of the cell type-specific marker genes were not detected in the proteomics analysis—almost certainly due to technical limitations. Our analyses support previous reports that NOD, AJ, and BALBc mice exhibit age-related neuronal loss (58, 59).

Our transcriptomic analyses identified age-related photoreceptor loss in WSB mice. Indeed, WSB mice exhibited hyperpigmented clumps on fundus exam, which was strikingly reminiscent of human RP phenotypes (76). Consistent with human RP, WSB mice had drastic photoreceptor loss with age. This loss appeared to be largely rod-specific, as scotopic ERG a-waves exhibited significant decline with age, and consequential inner retinal signaling was substantially blunted (as indicated by lower b-wave and nearly non-existent OPs). Patients with RP can exhibit severely reduced or absent OPs (77). Cone photoreceptors appeared to be subtly lost with age (as evident by a more subtle loss of photopic ERG amplitudes), and RGCs were relatively spared from this degeneration (as indicated by comparable RBPMS+ cell loss to B6 mice along with intact PERG responses). Interestingly, WSB photoreceptor loss was most severe at the central retina. This contrasts with RP disease progression—where peripheral rods are lost first—and is more consistent with patterns of photoreceptor loss in observed in human AMD. The central retina of the mouse shares some similarities to the human macula, including a higher density of photoreceptors, a thinner Bruch’s membrane, and fewer retinal pigment epithelium cells per photoreceptor (78). Thus, WSB mice model phenotypes relevant to multiple retinal degenerative diseases.

To investigate mechanisms relevant to photoreceptor degeneration in WSB mice, we compared the retinal proteome of 4M WSB mice to other pigmented strains. These results indicated that WSB mice exhibit DEPs involved in essential photoreceptor functions—including mitochondrial and retinol metabolism, and ciliary proteins within the BBSome (64-67, 79). These analyses predict the mechanisms that initiate the RP-like phenotype are intrinsic to photoreceptors. Importantly, WSB mice are not known to carry any *rd* mutations (60). Thus, WSB mice are likely to harbor novel mutations driving photoreceptor loss, which could be uncovered using genetic mapping studies. Altogether, these data suggest that WSB mice may be a unique model with which one could study mechanisms of age-related retinal degeneration relevant to human AMD and RP.

Our transcriptomic analyses also predicted an age-related loss of RGCs in NZO mice. Fundus examination of NZO mice revealed age-associated incidences of spots akin to cotton wool spots, exudates, and what appeared to be an epiretinal membrane in aged mice. Cotton wool spots and exudates are commonly found in many retinal vascular diseases, including DR (75, 80). These abnormalities are thought to arise from microvascular dysfunction and may indicate areas of local ischemia and blood-retina barrier breakdown (75, 80). Future work will be required to better understand whether the epiretinal membrane is similar to that observed in human patients. The presence of these abnormalities has not been previously reported in spontaneous mouse models of DR but cotton wool spots have been observed in a primate model of DR after chronic type 2 diabetes (81). In fact, widely used mouse models of DR fail to recapitulate the normal progression of disease pathology, and most models are limited to phenotypes associated with early stages of DR (81).

In DR, it is thought that initial hyperglycemia and metabolic dysfunction drives microvascular damage, promoting vascular compromise, eventually resulting in an increasingly hypoxic environment, promoting deleterious neovascularization (75, 81). The precise timing of neuronal loss amidst the microvascular dysfunction is not entirely clear in human patients, however, there are reports that there is loss of INL cells and RGCs even prior to overt DR (82). NZO mice are a polygenic model of metabolic syndrome exhibiting profound obesity, hypertension, hyperlipidemia, and insulin resistance, with approximately 50% of male mice becoming overtly hyperglycemic with age (69, 71). We observed no clear sex-dependent differences in the progression of NZO-associated retinal phenotypes, suggesting these changes may be due to the milieu of metabolic impairments NZO animals develop beyond hyperglycemia alone.

To probe the molecular causes of retinopathy in NZO mice, we investigated early retinal proteomic differences in NZO retinas. The retinal proteome of 4M NZO showed reduced expression of glutathione metabolic process and cellular detoxification pathways relative to other strains. In fact, the GSS protein (which is essential to produce glutathione) was significantly downregulated in NZO retinas relative to other strains. Glutathione levels are significantly reduced in many retinal diseases (83). Furthermore, levels of the glutathione-dependent enzyme GLO1 were significantly lower in 4M NZO retinas. GLO1 is a major detoxification enzyme for methylglyoxal, a precursor of advanced glycation end products (AGEs) (84, 85). Methylglyoxal, AGEs, and GLO1 have been linked to DR progression in patients and in rodent models (84-88). Altogether, reductions in these protective pathways likely increase susceptibility to oxidative stress-induced damage, which may be mediated by AGEs resulting from systemic metabolic dysfunction (89).

In accordance with the progression of human DR and retinal vascular disease, NZO animals displayed significant microvascular dysfunction relative to either WSB or B6 mice at 12M by multiple measures, including fluorescein angiography, FITC-Dextran leakage, and presence of acellular capillaries. NZO mice also exhibited age-associated loss of RBPMS+ RGCs and INL thickness, which manifested as reduced visual function as measured by both PERG and ERG. Collectively, these data suggest NZO mice better model the progression of human retinal vascular disease and DR than existing models, which includes the stepwise development of vascular insufficiencies and neuronal dysfunction.

In totality, our efforts: 1) generated a publicly available multi-omics database for retinal aging research across diverse genetic contexts to facilitate hypothesis development and refinement, 2) identified a common retinal aging signature across mice suggestive of reduced photoreceptor function, 3) determined that genetic context is a major driver of the molecular changes associated with retinal aging, 4) characterized WSB mice as a model for overt age-related and regionalized rod-specific photoreceptor loss, and 5) identified NZO mice as a novel mouse model of retinal vascular disease and RGC loss with striking similarities to human disease.

## Conclusion

Our study provides a valuable multi-omics resource for those investigating the aging retina, mechanisms of neurodegeneration, or the retina as a biomarker for changes within the brain. Our inclusion of the eight founder strains of the CC and DO diverse mouse populations provides important context for other phenotype- andexpression-based analyses conducted with these populations. From these data, we identified and validated two novel models of age-related retinal neurodegeneration.

## List of Abbreviations

ERG: Electroretinogram
PERG: Pattern Electroretinogram
RP: Retinitis pigmentosa
DR: Diabetic Retinopathy
AMD: Age-related Macular Degeneration
CC: Collaborative Cross
DO: Diversity Outbred
OCT: Optical Coherence Tomography
ONL: Outer Nuclear Layer
INL: Inner Nuclear Layer
RGC: Retinal Ganglion Cell
GCL: Ganglion Cell Layer
NFL: Nerve Fiber Layer
IPL: Inner Plexiform Layer
RBPMS: RNA Binding Protein, MRNA Processing Factor
PCA: Principal Component Analysis
ACAT1: Acetyl-CoA Acetyltransferase 1
OP: Oscillatory Potential
H&E: Hematoxylin and Eosin
GS: Glutathione synthetase
GLO1: Glyoxalase 1
AGEs: advanced glycation end products

## Declarations

**Availability of data and materials**

## Data availability

Bulk RNA-seq and proteomics data have been deposited at GEO and Proteome Exchange PRIDE respectively and are publicly available as of the date of publication. A shiny app for querying these data sets is available at: https://thejacksonlaboratory.shinyapps.io/Howell_AgingRetinaOmics/

## Author contributions

O.J.M., M.M., and G.R.H. designed the study. O.J.M., and M.M. developed experimental protocols, completed *in vivo* and IHC/histology experiments and analyses. M.M., O.J.M., T.L.C., A.A.H., C.A.D., and A.M.R. all euthanized mice and dissected tissues for various experiments. M.M. performed bioinformatics analyses. D.A.S. aligned RNA-seq data to strain-specific mouse genomes and contributed to study design. O.J.M., M.M, and G.R.H. wrote the manuscript. All authors approved the final version.

## Ethics approval and consent to participate

All research was approved by the Institutional Animal Care and Use Committee (IACUC) at The Jackson Laboratory.

## Consent for publication

Not applicable.

## Acknowledgments

We are grateful to Drs. Greg Carter, Patsy Nishina and Martin Pera for discussions on experimental design. We are also grateful to Dr. Kristen Onos, Kelly Keezer, and Melanie Maddocks-Goodrich for help in establishing and maintaining the strains for multi-omics analyses. We gratefully acknowledge the contribution of Mass Spectrometry and Protein Chemistry of the Protein Sciences, Genomic Technologies, Histology, and Microscopy Cores at The Jackson Laboratory for expert assistance with this publication. We thank Dr. Cara Hardy for her careful reading of the manuscript. We also would like to thank Dr. John Bachman for assisting with tissue collection and careful reading of the manuscript.

## Supplementary Materials

### This supplementary file includes

*Additional File 1. Supplemental Figure 1. Multi-omics profiling of the aging retina across genetically diverse mice reveals common molecular aging signatures*.

*Additional File 2. Supplemental Figure 2. Aged NZO mice develop epiretinal membranes*.

*Additional File 3. Supplemental Figure 3. WSB mice exhibit abnormal retinal structures by histology*.

*Additional File 4. Supplemental Figure 4: 4M WSB mice altered photoreceptor outer segment and metabolic DEPs relative to pigmented mice*.

*Additional File 5. Supplemental Figure 5. NZO mice develop microaneurysms and hemorrhages*.

*Additional File 6. Supplemental Figure 6: 4M NZO mice exhibit reduced expression of antioxidant pathway proteins relative to pigmented strains*.

*Additional File 7. Supplemental Table 1.* Excel file containing diagram of total numbers of mice used for transcriptomics and proteomics by strain, age, and sex.

*Additional File 8. Supplemental Table 2.* Excel sheets containing the common aging signature DEG and DEP analysis results, and the results of GO enrichment analyses for both the common aging signature DEGs and DEPs. For **DEG** analyses: *N*=8 (4F,4M) NZO, 129S1, and B6 mice at each age; NOD mice: *N*=8 (4F,4M) at 4M*, N*=7 (3F,4M) at 12M, *N*=6 (2F,4M) at 18M; AJ mice: *N*=8 (4F,4M) at 4M and 12M, *N*=7 (4F,3M) at 18M; BALBc mice: *N*=7 (3F,4M) at 4M and 12M, *N*=4F at 18 M; PWK mice: *N*=8 (4F,4M) at 4M and 12M, *N*=3F at 18M; CAST mice: *N*=8 (4F,4M) at 4M and 12M, *N=7* (3F,4M) at 18M; WSB mice: *N=7* (3F,4M) at 4M, *N*=8 (4F,4M) at 12M and 18M. For **DEP** analyses: *N*=8 (4F,4M) NZO, 129S1, and CAST mice at each age; NOD mice: *N*=7 (4F,3M) at 4M*, N*=8 (4F,4M) at 12M, *N*=6 (2F,4M) at 18M; AJ mice: *N*=8 (4F,4M) at 4M and 12M, *N*=7 (4F,3M) at 18M; BALBc mice: *N*=7 (3F,4M) at 4M, *N*=8 (4F,4M) at 12M, *N*=4F at 18 M; PWK mice: *N*=8 (4F,4M) at 4M and 12M, *N*=3F at 18M; B6 mice: *N*=8 (4F,4M) at 4M and 18M, *N=7* (3F,4M) at 12M; WSB mice: *N=7* (3F,4M) at 4M, *N*=7 (4F,3M) at 12M, *N*=8 (4F,4M) at 18M.

*Additional File 9. Supplemental Table 3.* Excel sheets containing the proteomics analysis of DEPs for 4M WSB vs the other pigmented strains, and the results of STRING-dB protein-protein interaction analysis and GO term enrichment. *N*=8 (4F,4M) NZO, 129S1, PWK, B6 and CAST mice; WSB mice: *N=7* (3F,4M).

*Additional File 10. Supplemental Table 4.* Excel sheets containing the proteomics analysis of DEPs for 4M NZO vs the other pigmented strains, and the results of STRING-dB protein-protein interaction analysis and GO term enrichment. *N*=8 (4F,4M) NZO, 129S1, PWK, B6 and CAST mice; WSB mice: *N=7* (3F,4M).

---

**Supplemental Figure 1.**
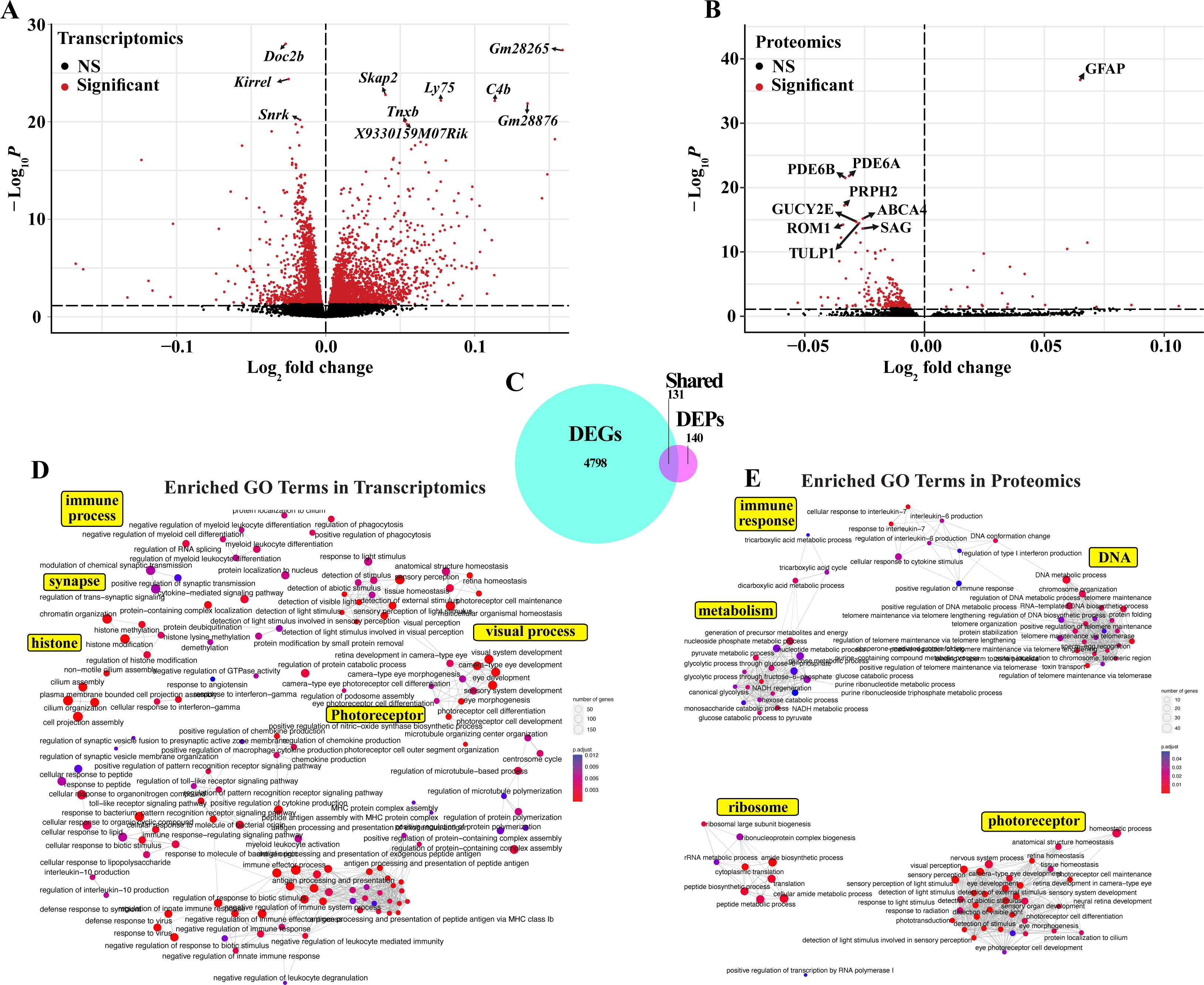
Multi-omics profiling of the aging retina across genetically diverse mice reveals common molecular aging signatures. Volcano plots of differentially expressed **A.** genes (DEGs) and **B.** proteins (DEPs) associated with the common retinal aging signature across strains. Genes and Proteins with FDRLJ<LJ0.05 are colored red. **C.** Venn diagram illustrating the overlap of DEGs and DEPs associated with the common aging signature across strains. Enrichment GO term plots of common aging signature **D.** DEGs and **E.** DEPs. Yellow boxes highlight a general theme of adjacent GO terms. In **A**,**D**: *N*=8 (4F,4M) NZO, 129S1, and B6 mice at each age; NOD mice: *N*=8 (4F,4M) at 4M*, N*=7 (3F,4M) at 12M, *N*=6 (2F,4M) at 18M; AJ mice: *N*=8 (4F,4M) at 4M and 12M, *N*=7 (4F,3M) at 18M; BALBc mice: *N*=7 (3F,4M) at 4M and 12M, *N*=4F at 18 M; PWK mice: *N*=8 (4F,4M) at 4M and 12M, *N*=3F at 18M; CAST mice: *N*=8 (4F,4M) at 4M and 12M, *N=7* (3F,4M) at 18M; WSB mice: *N=7* (3F,4M) at 4M, *N*=8 (4F,4M) at 12M and 18M. In **B,E**: *N*=8 (4F,4M) NZO, 129S1, and CAST mice at each age; NOD mice: *N*=7 (4F,3M) at 4M*, N*=8 (4F,4M) at 12M, *N*=6 (2F,4M) at 18M; AJ mice: *N*=8 (4F,4M) at 4M and 12M, *N*=7 (4F,3M) at 18M; BALBc mice: *N*=7 (3F,4M) at 4M, *N*=8 (4F,4M) at 12M, *N*=4F at 18 M; PWK mice: *N*=8 (4F,4M) at 4M and 12M, *N*=3F at 18M; B6 mice: *N*=8 (4F,4M) at 4M and 18M, *N=7* (3F,4M) at 12M; WSB mice: *N=7* (3F,4M) at 4M, *N*=7 (4F,3M) at 12M, *N*=8 (4F,4M) at 18M.

**Supplemental Figure 2.**
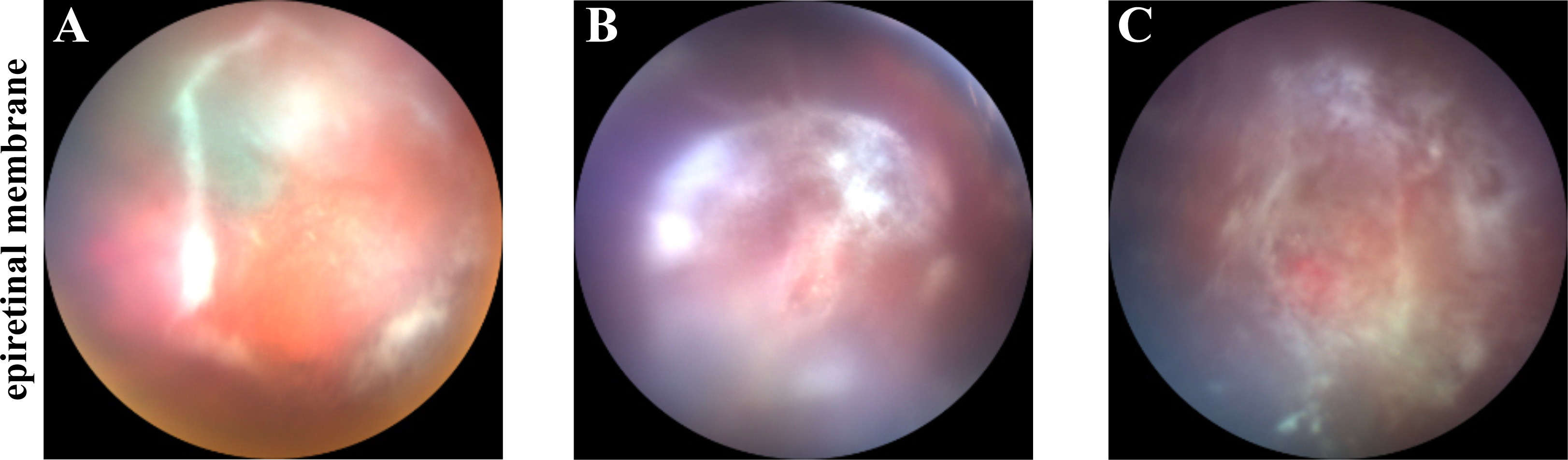
Aged NZO mice develop epiretinal membranes. **A-C.** Fundus images of epiretinal membranes we detected exclusively in 18M NZO mice.

**Supplemental Figure 3.**
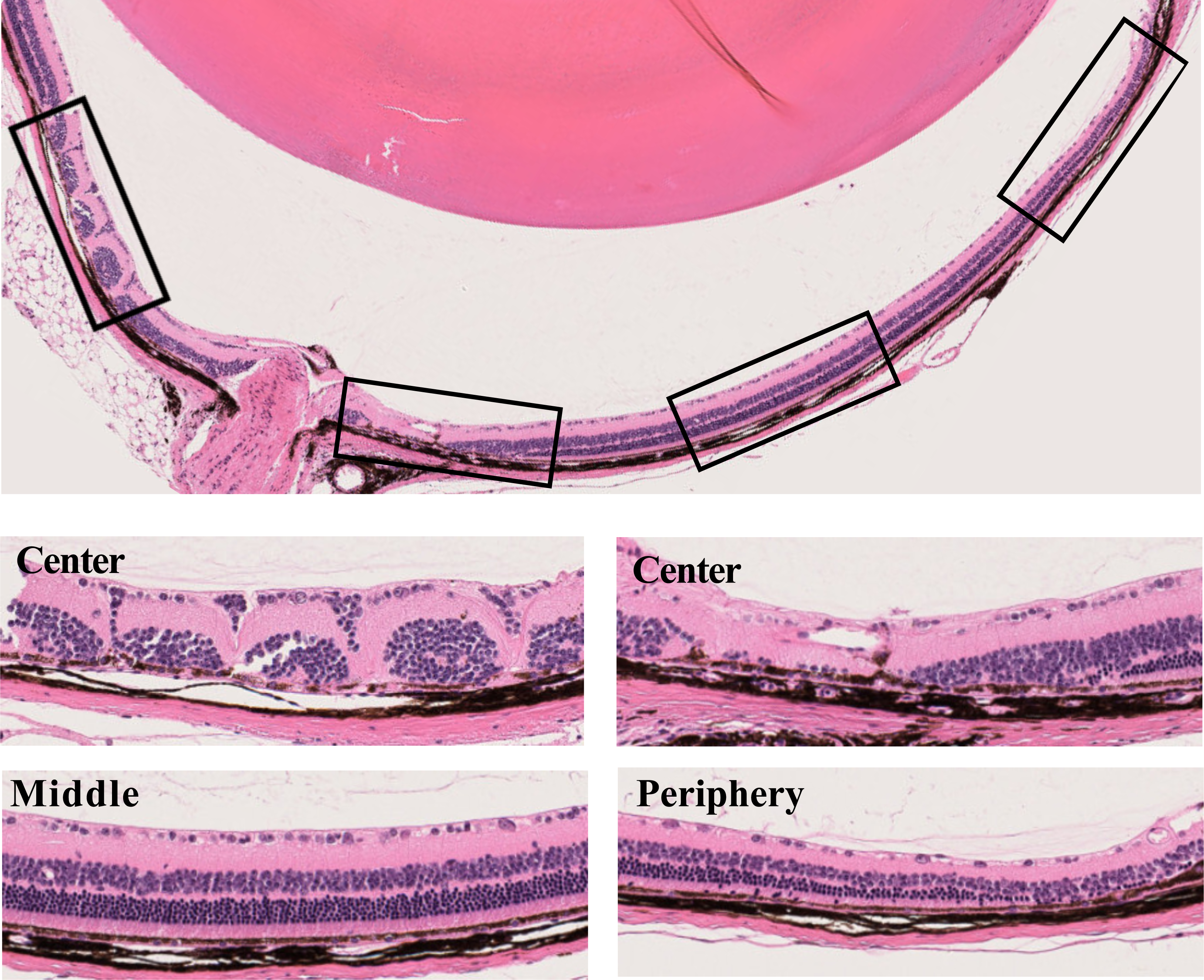
WSB mice exhibit abnormal retinal structures by histology. Representative images of abnormal retinal structures identified in aged WSB animals, which were never observed in young mice.

**Supplemental Figure 4:**
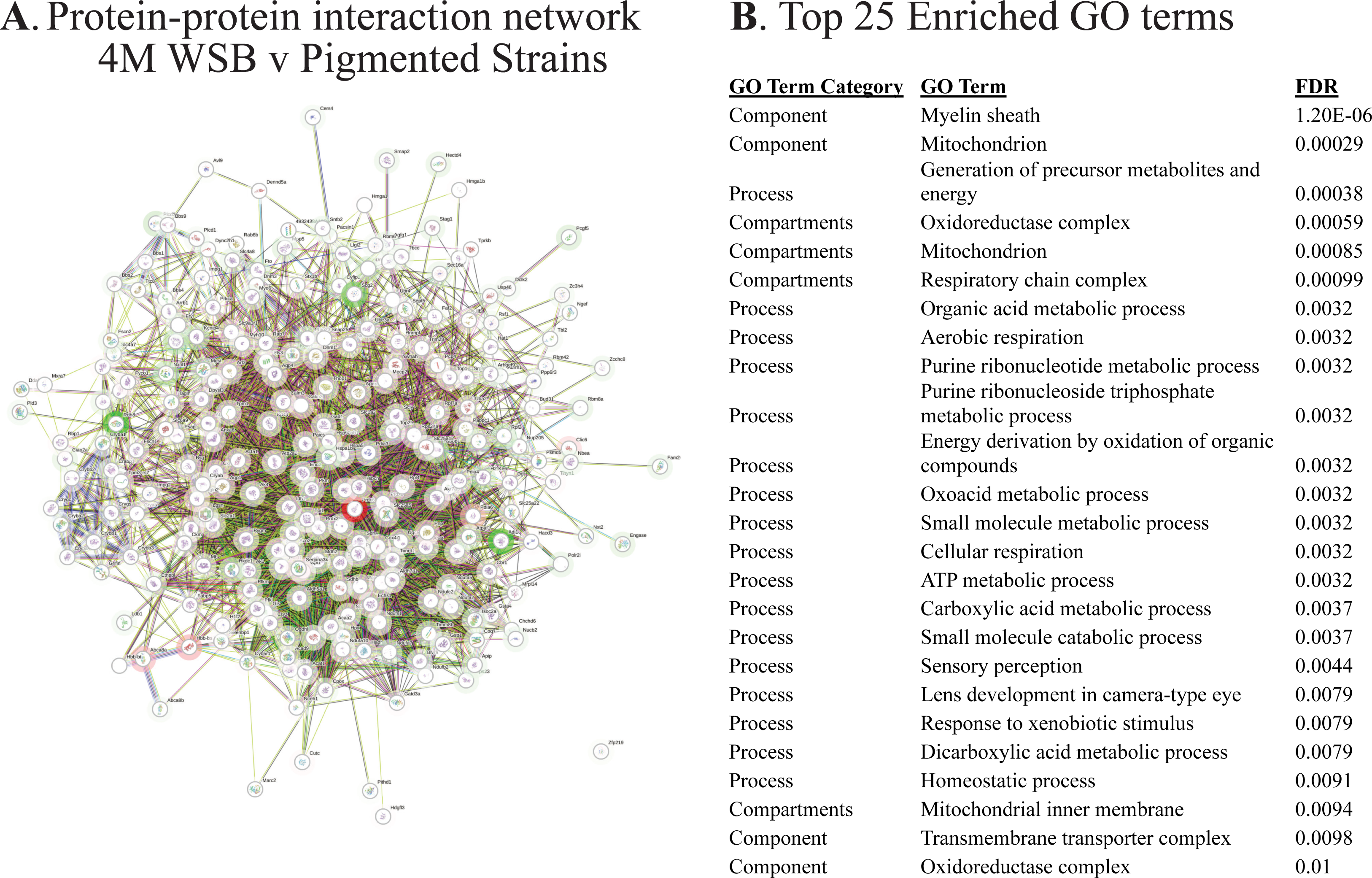
4M WSB mice altered photoreceptor outer segment and metabolic DEPs relative to pigmented mice. **A**. Physical STRING protein-protein interaction network of DEPs associated with 4M WSB vs other pigmented strains. 258 total proteins, 3247 interactions, *p*<0.0001. Node glow is colored by log_2_(fold-change) magnitude. Lines indicate physical interactions. **B**. Top 25 Enriched GO terms with FDR < 0.05 associated with the DEPs identified between 4M WSB and the other pigmented strains. *N*=8 (4F,4M) NZO, 129S1, PWK, B6 and CAST mice; WSB mice: *N=7* (3F,4M).

**Supplemental Figure 5.**
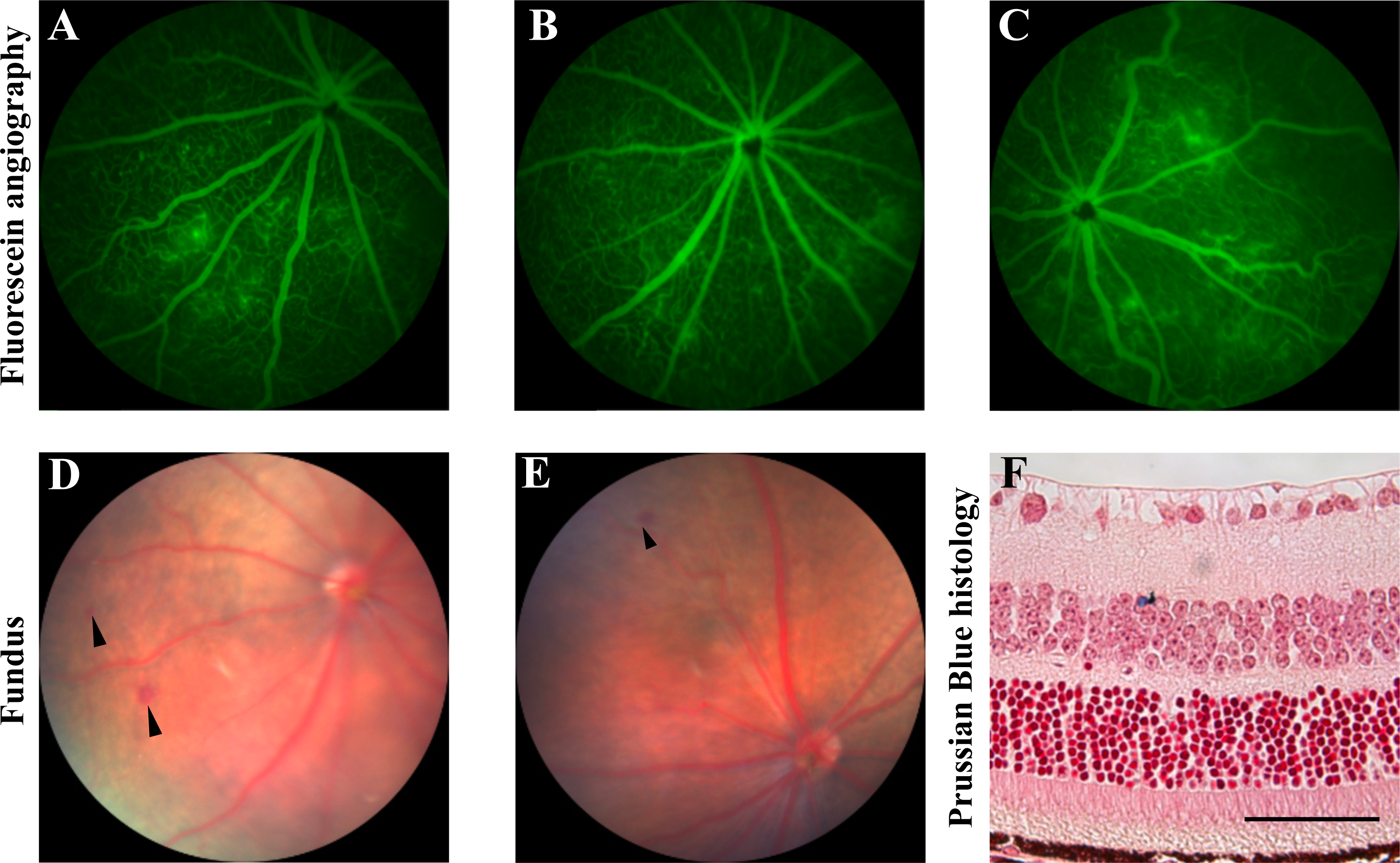
NZO mice develop microaneurysms and hemorrhages. **A-C.** Representative fluorescein angiography images of microaneurysms, and hemorrhages in 12-18M NZO mice. Notable vessel tortuosity is also visible. **D-E.** Fundus images of NZO mice showing deep red spots suggestive of hemorrhages. **F.** Prussian blue histological staining of an NZO retina indicating a hemorrhage event within the inner nuclear layer. Nuclei are stained pink with Fast Red. Scale bar: 50µm.

**Supplemental Figure 6:**
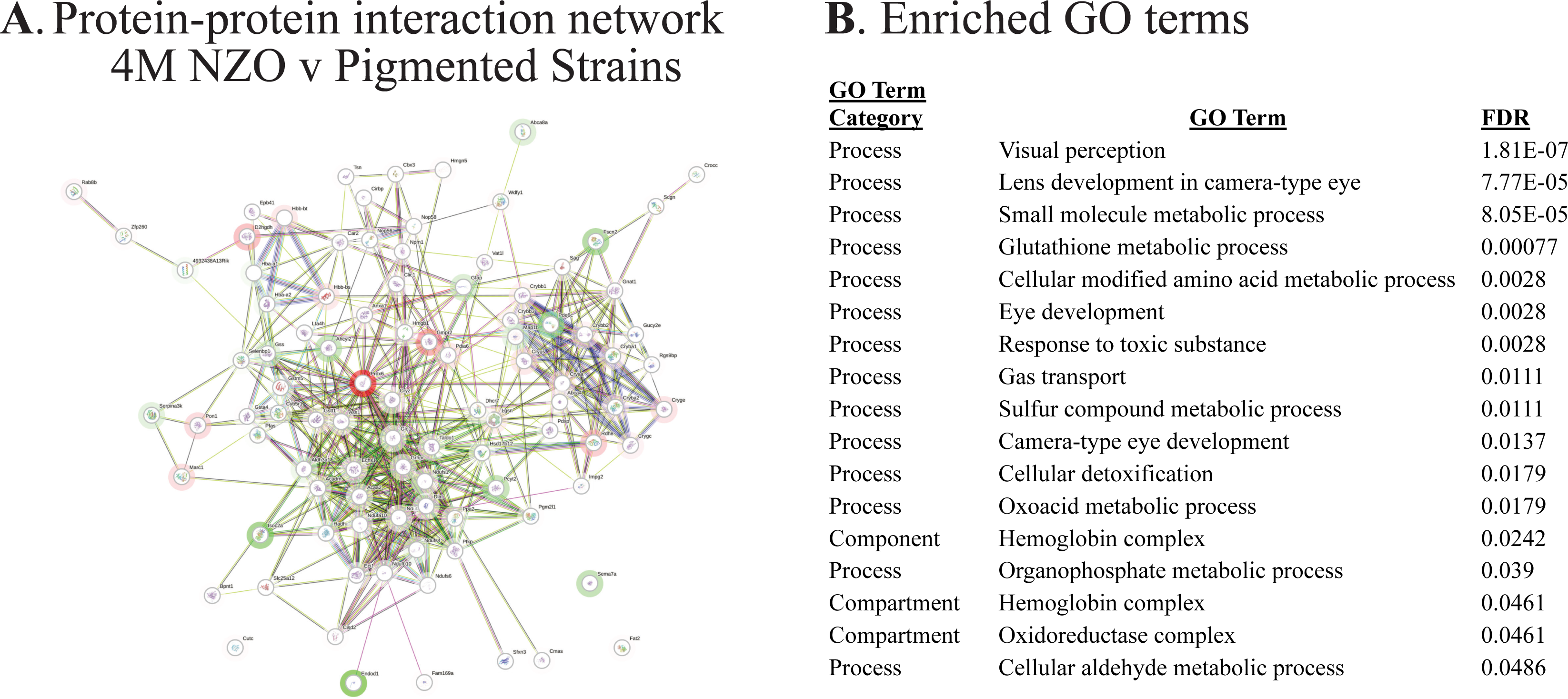
4M NZO mice exhibit reduced expression of antioxidant pathway proteins relative to pigmented strains. **A**. Physical STRING protein-protein interaction network of DEPs associated with 4M NZO vs other pigmented strains. 97 total proteins, 488 interactions, *p*<0.0001. Node glow is colored by log_2_(fold-change) magnitude. Lines indicate physical interactions. **B**. Enriched GO terms with FDR < 0.05 associated with the DEPs identified between 4M NZO and other pigmented strains. *N*=8 (4F,4M) NZO, 129S1, PWK, B6 and CAST mice; WSB mice: *N=7* (3F,4M).

